# TIM22 Complex-NADH dehydrogenase crosstalk maintains mitochondrial health by modulating cell death

**DOI:** 10.64898/2026.06.19.733344

**Authors:** Aishita Chakraborty, Rachayeeta Deb, SreeDivya Saladi, Patrick D’Silva

**Author notes:** Corresponding author: Patrick D’Silva.

## Abstract

Programmed cell death is essential for organismal development. When dysregulated, it leads to neurodegenerative diseases and cancer. Although apoptotic pathways are well studied, the role of mitochondrial import translocases in regulating cell death remains unclear. Our study reveals a unique apoptotic pathway controlled by the TIM22 complex, an inner mitochondrial membrane translocase. This pathway involves a multiprotein complex formed by Tim22 and Nde1, a part of the respiratory electron transport chain. Under stress, the cytosol-exposed Nde1 isoform, a pro-apoptotic factor, is stabilised by the TIM22 complex, which includes the Tim18 subunit and Tim22’s transmembrane segments. Notably, impairing the TIM22 complex and deleting Nde1 suppresses apoptosis and restores mitochondrial health. Beyond its role in import, our study uncovers a moonlighting function of the TIM22 complex in regulating mitochondria-dependent apoptotic cell death.

## Introduction

Mitochondria are double-membrane-bound dynamic organelles. They have diverse roles in energy production, macromolecular biosynthesis, cellular physiology, and metabolism (Harbauer *et al*, 2014). Most mitochondrial proteins are nuclear-encoded and imported via translocase machineries in the outer and inner mitochondrial membrane (Baker *et al*, 2007; Neupert & Herrmann, 2007; Wiedemann & Pfanner, 2017). After import, these proteins are compartmentalised to their destinations for proper mitochondrial functions (Kang *et al*, 2018; Kasahara & Scorrano, 2014; Nunnari & Suomalainen, 2012; Pfanner *et al*, 2019). If there is a defect in protein import, misfolded proteins accumulate, which then triggers stress signals (Palmer *et al*, 2021). As a consequence, the outer mitochondrial membrane becomes permeabilised, initiating a pro-apoptotic signaling cascade, that disrupts normal cell function (Dadsena *et al*, 2021). During this process, mitochondrial components such as cytochrome c, SMAC/DIABLO, Endo G, and AIF apoptotic factors are released into the cytosol, ultimately leading to cell death (Mustafa *et al*, 2024; Wang, 2001).

Intriguingly, another class of apoptogenic factor recently identified in yeast is NADH dehydrogenase Nde1, which is also a part of the respiratory chain complex (Herrmann & Riemer, 2021; Saladi *et al*, 2020). Nde1 and its paralogs, Nde2 and Ndi1, are non-proton pumping NADH dehydrogenases in yeast. These proteins transfer the electrons from NADH to ubiquinone. They act as a functional counterpart to mitochondrial respiratory complex I in higher eukaryotes (Luttik *et al*, 1998; Small & McAlister-Henn, 1998). Among these paralogs, Nde1 has a higher turnover rate compared to its paralogs. This is because Nde1 exists in two isoforms: one in the intermembrane space (IMS-Nde1), which performs the NADH dehydrogenase activity, and a cytosol-exposed form (Cyto-Nde1) with cytotoxic potential that can promote cell death. Cyto-Nde1 accumulates and is cleaved into two fragments depending on the cell’s mitochondrial potential and metabolic status: the mitochondrial fragment (f_mito_) and the cytosolic fragment (f_cyto_). The f_cyto_ fragment accumulates in the cytosol, triggering apoptosis. A high-throughput genetic screen also revealed that Nde1 toxicity may be suppressed mainly in the TIM22 complex mutant background (Saladi *et al*., 2020). Another recent finding suggests that the apoptogenic factor, Granzyme B, lacks the canonical mitochondrial targeting signal. It enters mitochondria through the Sam50-Tim22 channels to induce apoptosis (Chiusolo *et al*, 2017). These findings imply that the TIM22 complex might have a secondary function as a potential mediator of cell death apart from its canonical role of protein import.

The TIM22 complex (carrier translocase) is one of the inner membrane translocases. It imports multi-spanning membrane proteins with internal hydrophobic targeting sequences (Chacinska *et al*, 2009; Dudek *et al*, 2013; Rehling *et al*, 2004; Sirrenberg *et al*, 1996). In yeast, this complex (∼300 kDa) comprises membrane-bound (Tim22 dimer, Tim54, Tim18, and Sdh3) and peripheral (Tim9, Tim10, Tim12, Tim8, and Tim13) components. These facilitate the insertion of inner mitochondrial membrane (IMM) proteins (Koehler, 2004; Rehling *et al*, 2003; van der Laan *et al*, 2010). The core protein Tim22 exists as a dimer and forms the twin pore channel, while the Tim54 subunit docks small Tim proteins onto the core complex (Kovermann *et al*, 2002; Qi *et al*, 2020). The other two subunits, Tim18 and Sdh3, are essential for the complex assembly (Gebert *et al*, 2011; Kerscher *et al*, 2000; Koehler *et al*, 2000). The small Tim proteins, Tim9 and Tim10, form a soluble complex in the intermembrane space (IMS), with a portion of Tim9 associated with the TIM22 complex via Tim12 (Webb *et al*, 2006). Tim8 and Tim13, also import specific non-carrier proteins (Gebert *et al*, 2008; Gentle *et al*, 2007). The function of Tim22 is conserved across phylogeny. However, humans possess two additional subunits, AGK and Tim29, which maintain the integrity of the complex (Callegari *et al*, 2016; Kang *et al*, 2016; Kang *et al*, 2017). Mutations in the TIM22, AGK and TIM8A subunits can cause diseases such as mitochondrial myopathy and neurodegeneration. This is mostly due to impaired mitochondrial protein import, leading to oxidative stress and cell death (Kang *et al*, 2019; Kang *et al*., 2017; Roesch, 2002).

However, the mechanisms by which the carrier translocase regulates cell death remain elusive. To explore this, we used *Saccharomyces cerevisiae* as a model system. Our central question is: Does the TIM22 complex modulate cell death pathways independently or in concert with other mitochondrial protein complexes? We found that the functional TIM22 complex mediates apoptosis in response to apoptotic stimuli. It interacts with the mitochondrial fragment (f_mito_) of Cyto-Nde1. Notably, two transmembrane regions of the Tim22 core protein and the Tim18 subunit are key mediators of apoptosis. These components facilitate complex interactions and regulate apoptosis. Collectively, our findings suggest that a functional TIM22 complex promotes apoptosis. Apoptotic cell death is suppressed when the complex is impaired.

## Results

### The impaired TIM22 complex and Nde1 alleviate cell death induced by acetic acid

The absence of Tim18 (a subunit of the TIM22 import complex) was previously reported to confer resistance to arsenic-induced apoptosis. This suggests that it acts as a potential stress sensor in yeast (Du *et al*, 2007). To understand the role of the functional Tim22 complex in cell death, we first examined how Tim18 deletion (impaired TIM22 complex) affects cell death induced by acetic acid stress in yeast. Although yeast can utilise acetate as a carbon source, high concentrations of acetic acid can trigger apoptosis (Falcone & Mazzoni, 2016; Giannattasio *et al*, 2013; Guaragnella & Bettiga, 2021). Furthermore, Nde1 was identified as a pro-apoptotic factor, and its absence was observed to rescue cells from acetic acid-induced cell death (Saladi *et al*., 2020).

To understand the crosstalk between the TIM22 complex and Nde1, we generated yeast knockout strains Δ*tim18*, Δ*nde1*, and Δ*tim18*Δ*nde1* (Table **S1**). We then performed growth phenotypic analysis using gradient concentrations of acetic acid at 30°C (Figs. **1A** and **EV1A**). WT and the deletion strains showed no significant growth defects in 50 mM acetic acid compared to the untreated condition. However, at 70 mM, the WT strain displayed greater sensitivity than the knockout strains (Fig. **EV1A**). Upon further increasing the concentration to 90 mM, the WT strain exhibited a more severe growth defect than the Δ*tim18* strain, which also showed stress sensitivity. In contrast, Δ*nde1* cells showed slight growth defects. Strikingly, we observed marked growth restoration in the Δ*tim18*Δ*nde1* strain. This indicates an additive phenotypic effect of Tim18 and Nde1 in mitigating the toxicity of high acetic acid concentrations (Fig. **1A**). Taken together, our results led us to hypothesise that these proteins together might play an essential role in regulating cell death.

**Figure 1.**
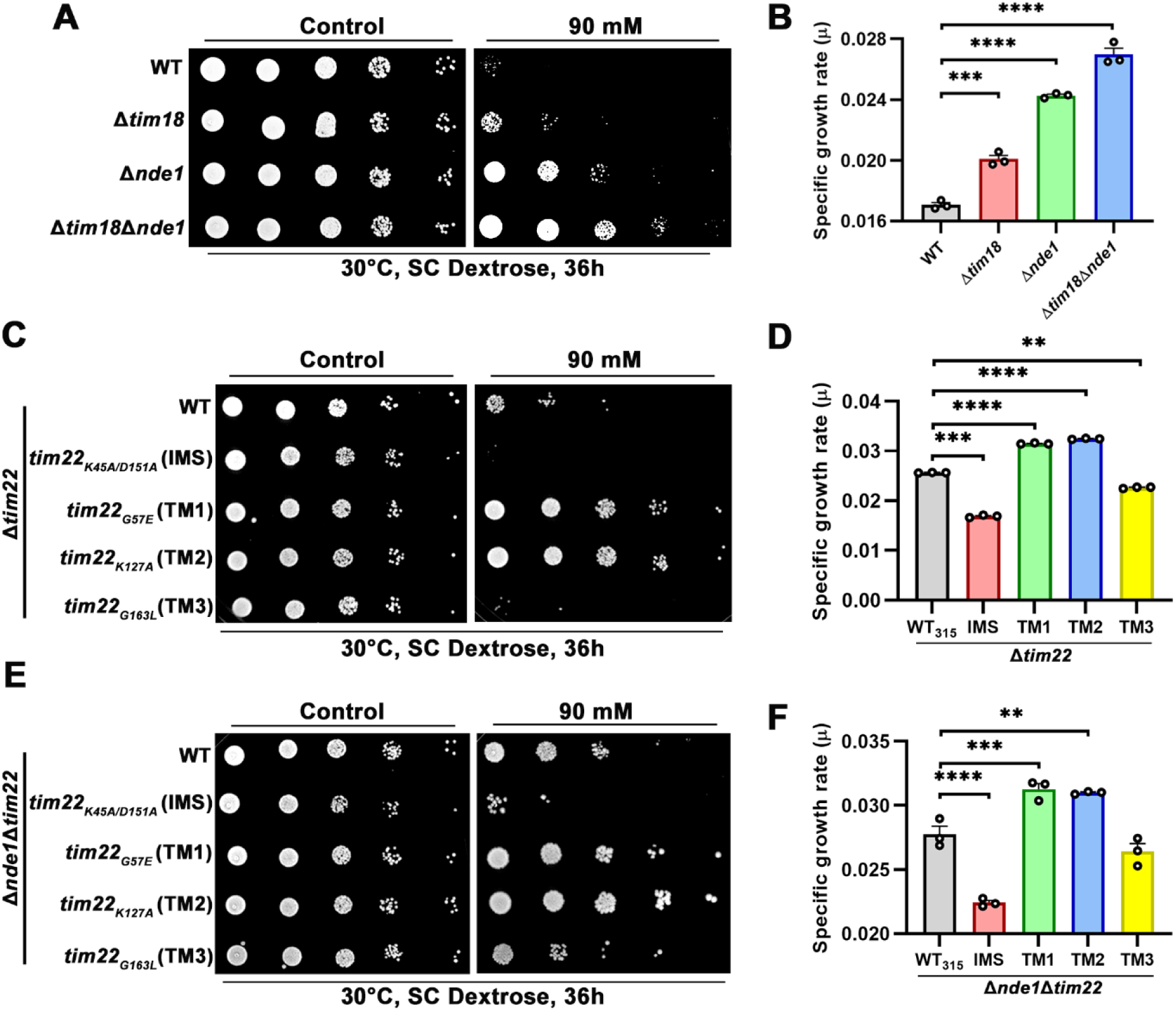
Acetic acid-mediated cell death via functional Tim22 complex requires Nde1. (**A**) Analysis of growth phenotype. WT, Δ*tim18,* Δ*nde1* and Δ*tim18*Δ*nde1* strains were treated with 90 mM acetic acid. Cells grown till mid-log phase in SC-Dextrose media at 30°C were exposed to 90 mM of acetic acid and incubated for 3h along with a set of untreated cells. Ten-fold serial dilutions of each sample were prepared and spotted on the indicated media. (**C, E**) *In vivo* phenotypic growth examination. Growth phenotype assessment in WT (where the *TIM22* gene has been expressed under pRS315 vector) and its IMS and TM mutants, as well as in absence of Nde1 was performed under 90 mM acetic acid. The strains were grown in SC-Dextrose till the mid-log phase and treated with the above-mentioned concentration of acetic acid for 3h. Both treated and untreated cells were then serially diluted tenfold and spotted on the indicated media. All images were acquired after 36 h of incubation at 30°C. (**B, D, and F**) The bar graphs represent the specific growth rates (µ) of the above-mentioned strains, calculated from their growth curves under acetic acid stress.

To validate our initial observations, we investigated dose-dependent effects of acetic acid in Tim22 mutant strains. These strains contained mutations in either the IMS or the three transmembrane domains (TM1, TM2 and TM3) in the Δ*tim22* background. We generated the Tim22 mutants using centromeric plasmids (see **Method Details** and Table **S1**). Growth phenotypic analysis was performed at 30°C (Kumar *et al*, 2020). At 50 mM acetic acid, we observed no significant difference in growth (Fig. **EV1B**). At 70 mM, WT_315_ (pRS315-*TIM22*), the IMS mutant (*tim22_K45A/D151A_*) and the TM3 mutant (*tim22_G163L_*) exhibited growth defects (Fig. **EV1B**), highlighting sensitivity. TM1 (*tim22_G57E_*) and TM2 (*tim22_K127A_*) mutants, however, exhibited robust growth rescue at 90 mM acetic acid (Fig. **1C**), confirming their resistance. These findings align with the accelerated growth observed in Tim18 and Nde1 deletion conditions, as TM1 and TM2 mutations are known to reduce Tim18 association with the core complex. Meanwhile, the high sensitivity of IMS and TM3 mutants at both 70 and 90 mM was likely due to impaired disulfide bond formation and defective translocase activity, respectively (Fig. **1C** and **EV1B**).

To further confirm the role of Tim22 and Nde1 in cell death, we deleted Nde1 in WT and Tim22 mutant backgrounds (Table **S1**). Phenotypic analysis under 90 mM acetic acid stress revealed enhanced growth in TM1 and TM2 mutants (Fig. **1E**). This suggests that these domains attenuate cell death more effectively in the absence of Nde1. This is consistent with the additive rescue effect observed in the Tim18 and Nde1 double deletion strain (Fig. **1A**). Meanwhile, the WT_315_, IMS, and TM3 mutants showed a mild growth phenotype in the Nde1 deletion background (Fig. **1E**). The specific growth rates of the respective strains under stress conditions, calculated from growth kinetics, correlated with the growth patterns observed in spot test analysis (Figs. **1B, 1D and 1F).**

To determine whether other components of the TIM22 complex contribute to cell death regulation, we generated an Sdh3 knockout strain (Table **S1)**. Growth-phenotype analysis under acetic acid showed that Sdh3 deletion did not confer resistance (Fig. **EV1C**). This suggests that Sdh3 plays a distinct role from Tim18 or Tim22. To test TIM22 complex specificity in cell death, we also deleted Tim21, a transmembrane subunit of the TIM23 complex (another prominent import complex of the IMM) (Chacinska *et al*, 2005; Mokranjac *et al*, 2005). Deletion of Tim21 also failed to resist acetic acid-induced cell death (Fig. **EV1D**). These findings indicate that only the functional TIM22 complex, together with Nde1, specifically regulates cell death.

Yeast mitochondria have three NADH dehydrogenase paralogs: Nde1, Nde2 and Ndi1 (Grandier-Vazeille *et al*, 2001). To clarify their involvement in programmed cell death, we generated combinatorial knockout strains of Nde2 and Ndi1 in WT and Δ*tim18* backgrounds (Table **S1**). Deleting Ndi1 or Nde2 alone did not rescue growth defects under 90 mM acetic acid stress. However, Δ*tim18*Δ*ndi1* and Δ*tim18*Δ*nde2* double deletion strains exhibited mild growth restoration, likely reflecting rescue from Tim18 deletion, as observed in Fig. **1A** (Figs. **EV1E** and **EV1F**). Overall, all these findings reinforce the specific role of the Tim22 complex in modulating cell death along with Nde1.

### The TIM22 complex and Nde1 synergistically induce cell death through apoptosis

Yeast cells undergo acetic acid-induced programmed cell death, which exhibits hallmarks similar to apoptosis in higher eukaryotes (Ludovico *et al*, 2001). In yeast, under acetic acid stress, Nde1 predominantly exists in its cytosolic isoform (Cyto-Nde1), which triggers apoptosis (Saladi *et al*., 2020). As acetic acid is a known apoptotic stress inducer (Guaragnella & Bettiga, 2021), we next sought to determine whether cell death mediated by the TIM22 complex and Nde1 proceeds through a common apoptotic pathway. A notable hallmark of apoptosis in yeast is the externalisation of phosphatidylserine to the outer leaflet of the plasma membrane and loss of membrane integrity, which can be detected by staining together with FITC-labelled Annexin V and PI (Bankapalli *et al*, 2020; Li *et al*, 2006; Madeo *et al*, 2009). Flow cytometry analysis revealed that the majority of WT_(genomic)_ and WT_315_ cells treated with 90 mM acetic acid were in the late apoptotic stage accounting for 94% and 91.7% of the cell population, respectively (Figs. **2A, 2C** and **2B, 2D**) . However, the percentage of late apoptotic cells reduced significantly in Δ*tim18* (67.1%) and Δ*nde1* (62.8%) strains. Surprisingly, a pronounced shift in the cell population from the late apoptotic to the early apoptotic stage was observed in Δ*tim18*Δ*nde1* strain (49.7%), as depicted in the dot plots and bar graphs (Figs. **2A** and **2C**). Similarly, we observed that the deletion of Nde1 in the WT_315_ background resulted in 69.8% late apoptotic cell population, which was comparable to that of the Δ*nde1* strain (62.8%). Interestingly, the Δ*nde1*/TM1 (45.3%) and Δ*nde1*/TM2 (41.5%) mutants displayed apoptotic profiles similar to the Δ*tim18*Δ*nde1* strain (49.7%) (Figs. **2B** and **2D**). Thus, these results clearly indicated the presence of a functional TIM22 complex, together with Nde1, promotes apoptotic cell death.

**Figure 2.**
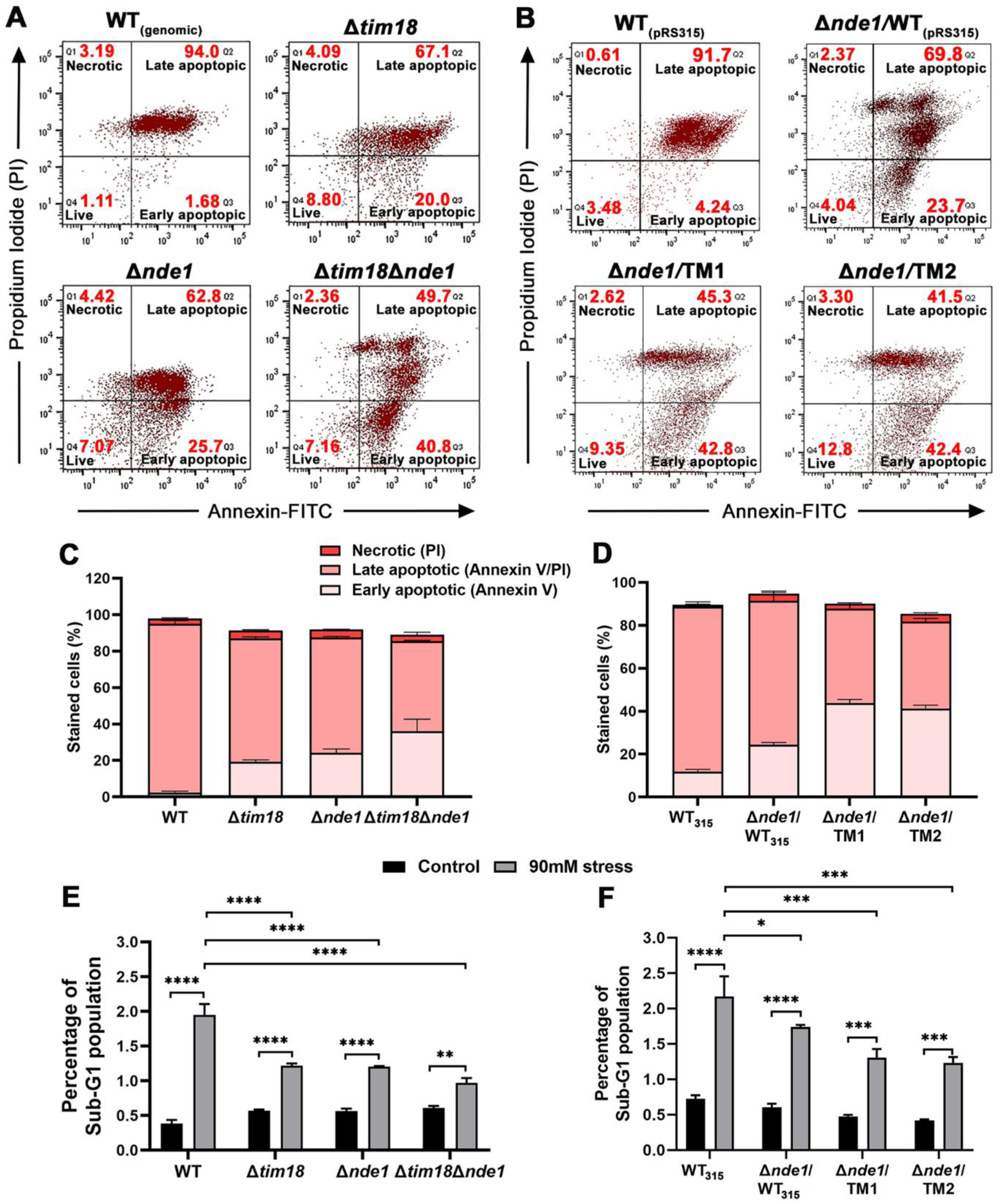
Apoptotic cell death analysis of the impaired carrier translocase machinery. (**A, B**) Apoptotic assay. All the studied strains were grown in SC-Dextrose until mid-log, then treated with 90 mM acetic acid, and stained with Annexin-V/PI for flow cytometry analysis. (**C, D**) The percentage of cells in different stages of apoptosis was quantified from three independent experiments (n=3) and represented as bar graphs. In each experiment, 30,000 cells were analysed. Error bars indicate n=3 biological replicates. (**E, F**) DNA content distribution under acetic acid stress. The studied strains were cultured in dextrose media and exposed to 90 mM stress. DNA was stained with propidium iodide and analysed by flow cytometry. The bar graphs represent the percentage of the sub-G1 population for the indicated strains under control and stress conditions. Quantification data were obtained from three independent experiments (n = 3). In each experiment, 30000 cells were analysed. Values of *p* ≤ 0.05 were considered significant (*), *p* ≤ 0.01 very significant (**), *p* ≤ 0.001 extremely significant (***) and *p* ≤ 0.0001 very extremely significant (****).

A fundamental correlation between apoptosis and DNA fragmentation has been reported (Del Carratore *et al*, 2002). To validate our hypothesis, we examined DNA content profiles of deletion and mutant strains under acetic acid treatment. Apoptotic cells were stained with PI and analysed by flow cytometry, revealing a distinct hypodiploid or sub-G1 peak, easily distinguished from the normal (diploid) G1 and G2/M peaks of the cell cycle. The WT and WT_315_ strains displayed significant sub-G1 populations, at 2.06% and 2.37%, respectively (Figs. **2E**, **2F** and **EV2A, EV2B)**. In contrast, the double deletion strain (Δ*tim18*Δ*nde1*), as well as the Δ*nde1* strain expressing either TM1 or TM2 mutant constructs (Δ*nde1*/TM1 and Δ*nde1*/TM2) demonstrated a marked reduction in the sub-G1 population (Figs. **2E**, **2F** and **EV2A and EV2B**).

Another characteristic marker of apoptosis is chromatin condensation, typically observed as condensed chromatin near the nuclear envelope (Clifford *et al*, 1996; Li *et al*., 2006). Fluorescence microscopy analysis revealed that after acetic acid treatment, the WT and WT_315_ cells exhibited significant DNA condensation, as indicated by DAPI staining (Figs. **EV2C** and **EV2D**). In contrast, the Δ*tim18*Δ*nde1* strain and TIM22 loss-of-function (TM1 and TM2) mutants lacking Nde1 displayed a considerable reduction in chromatin condensation (Figs. **EV2C** and **EV2D**). Quantitative analysis of intensity profiles (Figs. **EV2E** and **EV2G**) and corrected total cell fluorescence (CTCF) (Figs. **EV2F** and **EV2H**) showed similar results. Consistent with the cell viability results, it was observed that deletion of Tim18 or the absence of Nde1 in TM1 and TM2 mutants delayed the progression of apoptosis, thereby highlighting the importance of a functional TIM22 complex in the Nde1-dependent apoptotic pathway.

### Genetic and proteomic analysis identifies Nde1 as a functional partner of the TIM22 complex

Cellular protein synthesis and abundance are precisely controlled through multiple regulatory pathways. However, dysregulation of these processes can lead to protein accumulation, resulting in cytotoxicity and ultimately cell death (Kumar *et al*, 2023; Wrobel *et al*, 2015). Elevated levels of these mitochondrial precursor proteins can compromise the efficiency of the mitochondrial protein import machinery (Lionaki *et al*, 2023). Although the import of Nde1 depends on the presequence translocase or the TIM23 complex (Kizmaz *et al*, 2024), a high throughput screen suggested that mutants of the TIM22 complex conferred resistance to Nde1-mediated cell death (Saladi *et al*., 2020). These observations further prompted us to investigate the putative genetic interaction between Nde1 and the TIM22 complex. To rule out the possibility that insufficient Nde1 levels contributed to the observed phenotypes in the aforementioned deletion and mutant strains, we overexpressed *NDE1* gene in the pRS416_TEF_ vector under high-expression promoters (Table **S1**). Subsequently, growth phenotype and protein expression were analysed in these cells. Strikingly, overexpression of Nde1 in the WT strain showed a significant growth sensitivity. On the contrary, the Δ*tim18* strain did not exhibit any growth defect despite Nde1 overexpression (Fig. **3A**). Similarly, the WT_315_ and TM3 strains displayed growth defects under Nde1 overexpression, while the IMS mutant showed enhanced sensitivity. However, the TM1 and TM2 mutants were resistant to Nde1-mediated cell death (Fig. **3C**). These results were consistent with those observed under 90 mM acetic acid stress (Fig. **1**).

**Figure 3.**
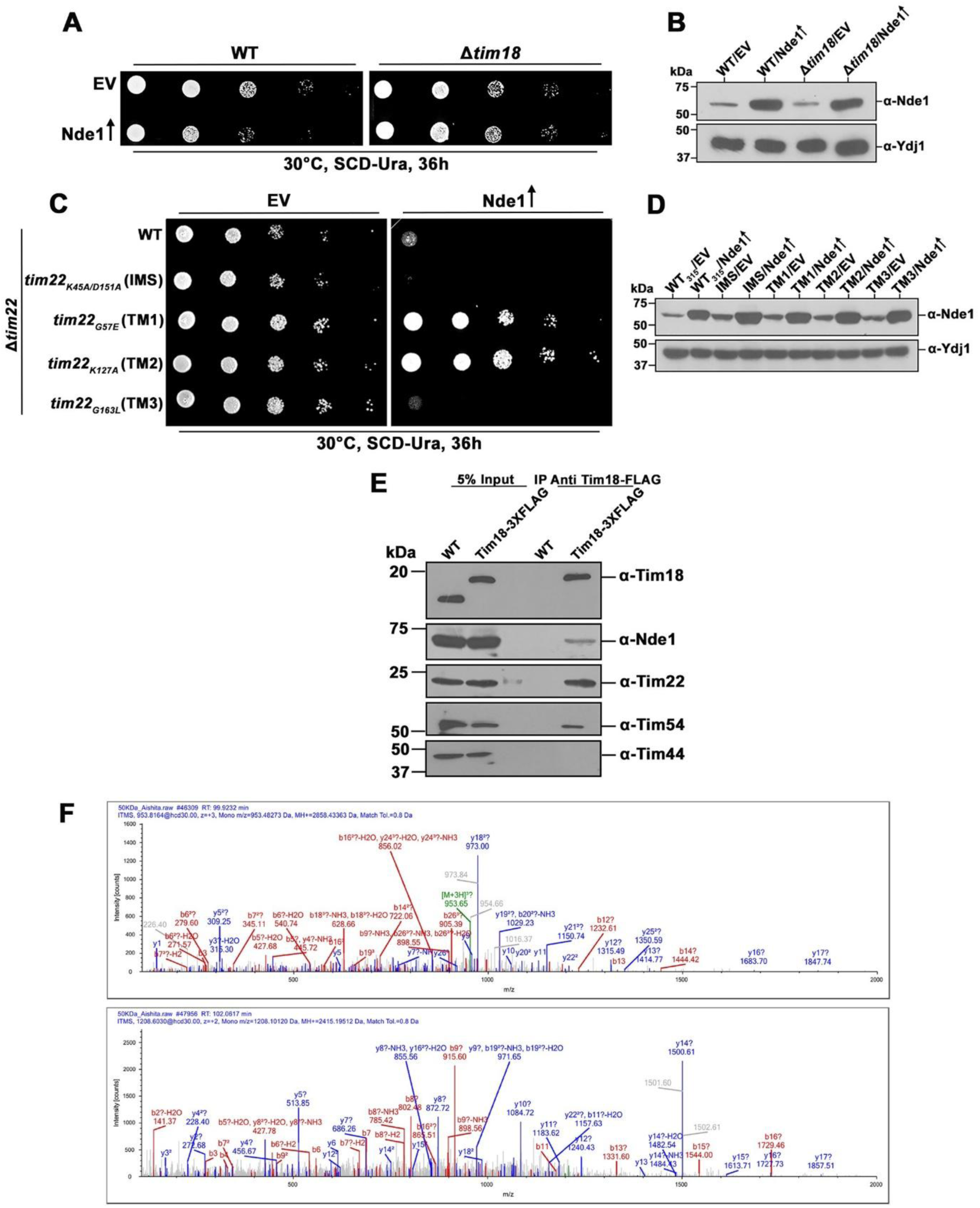
Nde1 lethality is ameliorated in the dysfunctional Tim22 complex, while its interaction is stabilised under a functional complex. (**A, C**) Growth phenotype examination upon overexpression of Nde1. WT and Δ*tim18* strains, as well as WT315 and its mutants overexpressing Nde1 were serially spotted on the indicated media at a permissive temperature overexpressing Nde1 under the control of centromeric plasmid pRS416TEF, were serially diluted and spotted on the indicated media at a permissive temperature. (**B, D**) Confirmation of Nde1 overexpression by Western blotting. The overexpression of Nde1 was measured by western blotting in whole-cell extracts of the indicated strains, where EV denotes the empty vector, and a vertical arrow indicates overexpression. (**C**) Co-IP assay to assess the interaction between Nde1 and the TIM22 complex. Mitochondrial isolates from the WT strain expressing Tim18-FLAG were subjected to Co-IP using anti-FLAG conjugated Protein G Sepharose beads. Bound proteins were detected by immunoblotting using specific antibodies. 5% of the total lysate (input) was used as a loading control. (**D**) The same co-immunoprecipitated sample was subjected to trypsin digestion followed by Orbitrap LC-MS analysis to identify the Nde1 binding peptide spectrum.

Overexpression of Nde1 in the deletion and mutant strains was confirmed by western blotting with an Nde1 antibody. Anti-Ydj1 served as the loading control (Figs. **3B** and **3D**). Thus, Nde1 overexpression induced cellular toxicity in strains with an intact TIM22 complex (WT and WT_315_), which suggests a strong genetic interaction between the proteins. Intriguingly, mutations in the TM1 and TM2 domains, characterised by Tim18 dissociation from the complex, showed tolerance to the Nde1-induced cytotoxicity, suggesting the existence of a compensatory mechanism associated with an impaired TIM22 complex.

Till now, our data indicated that the TIM22 complex, particularly the Tim18 subunit, is required for Nde1-mediated cell death. Next, we aimed to establish whether the overexpression of Tim18 alone, as a part of the functional TIM22 complex, could also induce cytotoxicity. To address this question, we overexpressed Tim18 in the WT and Δ*nde1* strains and assessed their growth phenotype. However, Tim18 overexpression had no significant effect on cellular growth in both strains, suggesting that Nde1 is the primary driver of cytotoxicity rather than Tim18 or the core Tim22 protein, and that Nde1’s apoptotic function depends on the presence of a functional TIM22 complex (Figs. **EV3A** and **EV3B**).

Furthermore, to validate the qualitative interaction between Nde1 and the TIM22 complex, coimmunoprecipitation coupled with mass spectrometry analysis was performed in WT untagged WT cells were used as control to exclude non-specific interactions. Mitochondrial lysates isolated from these strains were solubilised with digitonin buffer and incubated overnight with preconjugated anti-FLAG antibody. The immunoprecipitated protein samples were then analysed using western blotting and mass spectrometry. Strikingly, the Nde1 protein was detected as a distinct band migrating at ∼57 kDa region. In addition, Tim18 efficiently co-immunoprecipitated with Tim22 and Tim54 proteins, confirming the integrity of the TIM22 complex in the FLAG-tagged WT strain. Tim44, a component of the TIM23 complex, was not detected in the immunoprecipitants (Fig. **3E**), demonstrating the specificity of interaction between Nde1 and the TIM22 complex. To further characterise the interactome, the remaining immunoprecipitated sample was subjected to Orbitrap LC-MS analysis. Protein identification using Proteome Discoverer 2.5 software confirmed the presence of the Nde1 protein profile within the Tim18-FLAG interactome, along with the other interacting proteins (Fig. **EV3C**). MALDI-TOF mass spectra analysis of Nde1 was represented, validating with our Co-IP data (Fig. **3F**). The experimental workflow for the mass spectrometry is outlined in Fig. **EV3C**, followed by a proposed model detailing the transient interaction between the TIM22 complex and Nde1. Collectively, these findings indicate a physical and functional association between Nde1 and the TIM22 complex in the absence of stress.

### The functional TIM22 complex facilitates the formation of pro-apoptotic cytosolic Nde1

Based on the observed interaction between the TIM22 complex and Nde1 under physiological conditions, we further aimed to elucidate whether Nde1 associates with the functional TIM22 complex in a stress-dependent manner to facilitate cell death activity. Thus, we hypothesized that in order to mediate cell death together, Nde1 should form a supercomplex as it lies in proximity to the functional carrier translocase. Therefore, we performed BN-PAGE analysis, followed by western blotting in mitochondrial lysates isolated from deletion and mutant strains under both control and acetic acid-treated conditions. In wild-type cells, the TIM22 complex assembly was detected as a stable 300 kDa band under both control and stress conditions. A similar migration pattern was observed in the Δ*nde1* strain and the TM1 and TM2 mutants. In contrast, disruption of the complex was observed in Δ*tim18* strains, and in IMS and TM3 mutants, regardless of treatment (Figs. **4A** and **4C**). Furthermore, immunoblotting with an anti-Nde1 antibody revealed a higher order complex migrating at approximately 360 kDa in WT and TM2 strains, most likely representing the association of Nde1 with the core TIM22 machinery. However, only minor subcomplexes of this Nde1-TIM22 assembly were detected in the IMS and TM1 but the effect was more pronounced in Δ*tim18* and TM3 strains under control conditions, indicating partial destabilisation of the interaction. Notably, exposure to acetic acid stress led to a significant disruption of this interaction, resulting in complete dissociation of Nde1 from the supercomplex across all the studied strains (Fig. **4B** and **4D**). This suggests that the stability of Nde1, and specifically the interplay between its dual isoforms, may serve as a regulatory switch for cell death pathways. To assess the specificity of this interaction, the porin complex was analysed as a control, whose stability remained unaltered even under stress conditions (Fig. **4E** and **4F**). These findings indicate that the (IMS) Nde1-TIM22 interaction is a dynamic assembly that responds to mitochondrial physiological state (Fig. **4G**), necessitating further biochemical studies to elucidate its role in programmed cell death. Moreover, we checked the steady-state expression levels of Nde1 and Tim22 subunits under physiological conditions. To which we observed no significant alteration in the protein levels in the deletion and mutant strains. However, a mild reduction in the expression of Tim22 and Tim54 was observed in Δ*tim18* and IMS mutants, indicating their role in maintaining TIM22 complex integrity (Figs. **EV4**).

**Figure 4.**
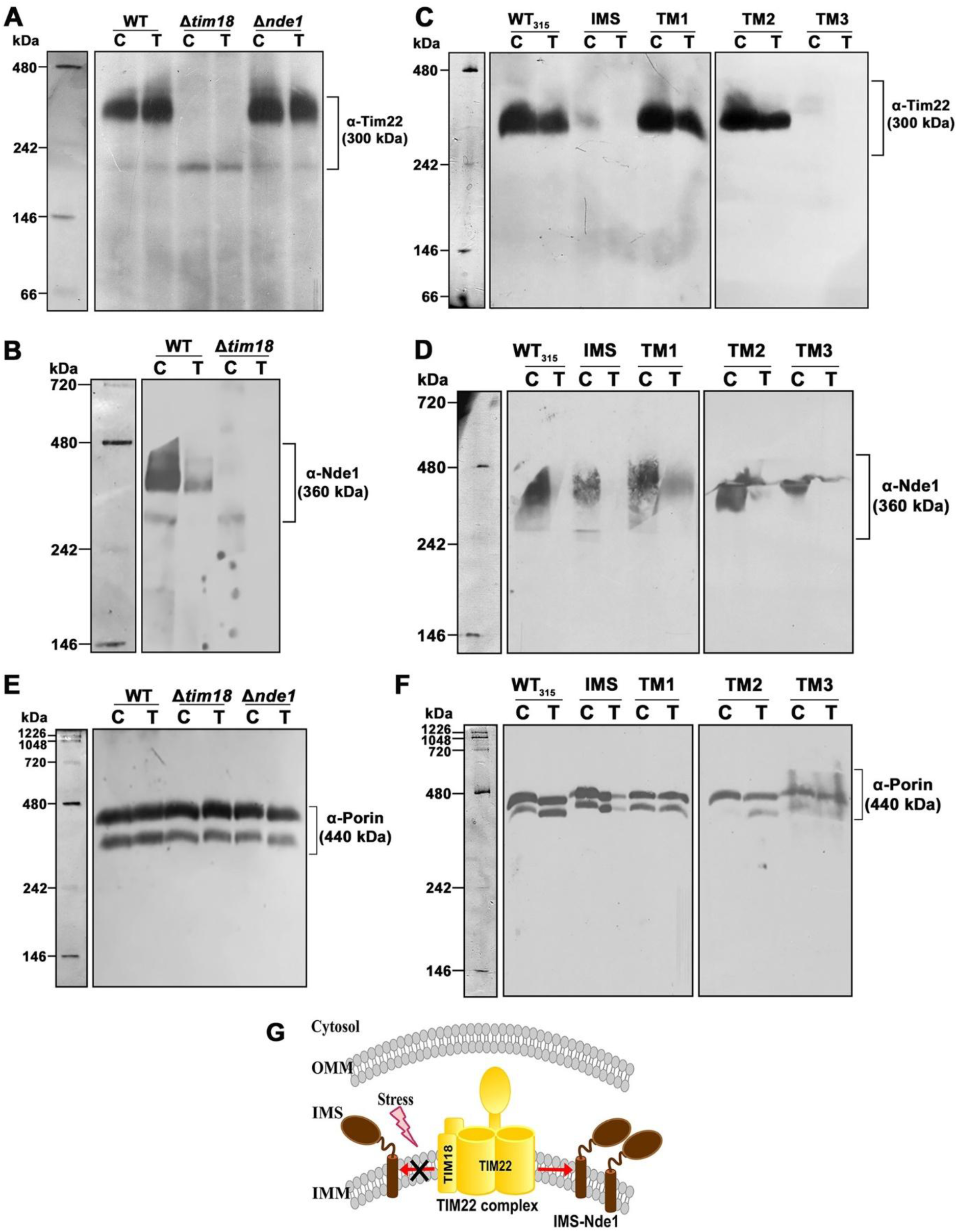
Nde1 forms a complex with the carrier translocase. (**A, C**) Stability of the Tim22 complex was analysed using BN-PAGE. Mitochondrial lysates isolated from WT and Δ*tim18* strains, and from the WT315, IMS, TM1, TM2, and TM3 strains under control and 90 mM acetic acid stress, were solubilised in digitonin buffer. The levels of Tim22 were subsequently examined using BN-PAGE followed by immunoblotting. (**B, D**) The stability of Nde1 complex formation with carrier translocase was analysed using BN-PAGE under normal and stress conditions. Mitochondria from the deletion and mutant strains were isolated and solubilised using a digitonin buffer. The levels of Nde1 were then analysed by immunoblotting. (**E, F**) Analysis of the porin complex from mitochondrial lysates was performed as a control to evaluate its stability under both treated and untreated conditions. **C** and **T** denote untreated and acetic acid-treated conditions, respectively. (**G**) Schematic representation of the IMS-Nde1 forming supercomplex with the Tim22 complex under normal conditions, but not under acetic acid stress.

However, the question remains open as to the possible mechanistic insights into how the complex formation of IMS-Nde1 with the TIM22 complex affects cell death activity. Previous studies reported that the f_cyto_ fragment of Cyto-Nde1, released into the cytosol, possesses cytotoxic potential, whereas the f_mito_ fragment of Cyto-Nde1 represents the mitochondrial remnant of the processed Cyto-Nde1 (Fig. **5I**) (Saladi *et al*., 2020). Thus, under stress conditions, expression levels of both the IMS-Nde1 (∼57 kDa) and the f_mito_ fragment (∼20 kDa) of the Cyto-Nde1 isoform were analysed in mitochondrial lysates isolated from both the deletion and mutant strains of the TIM22 complex. Interestingly, compared to control conditions, the WT and WT_315_ strains showed an increase in the f_mito_ fragment under stress (Figs. **5A** and **5E)**. However, a significant reduction in the levels of this fragment was observed in the Δ*tim18* strain (Figs. **5B**). IMS and TM3 strains displayed results similar to WT (Fig. **5E**). Interestingly, the f_mito_ levels decreased significantly in the TM1 and TM2 mutants, indicating a domain-specific role for the TIM22 complex in stabilising the Cyto-Nde1 isoform (Fig. **5E** and **5F**). Despite the complex instability in IMS and TM3 mutants, the presence of functional TM1 and TM2 domains helps these strains to retain the Cyto-Nde1 isoform via stabilisation of the f_mito_ fragment (Fig. **5E**). Interestingly, under stress, the levels of Tim22 and Tim18 remained stable in both WT and WT_315_, with a minor decline in Tim54 levels. Moreover, the loss of Nde1 did not significantly affect the protein levels of Tim22, Tim18 or Tim54, even under stress conditions, in the deletion strains (Figs. **5A** and **5E**). This result indicated that Nde1 stability during apoptosis depends on the functional TIM22 complex.

**Figure 5.**
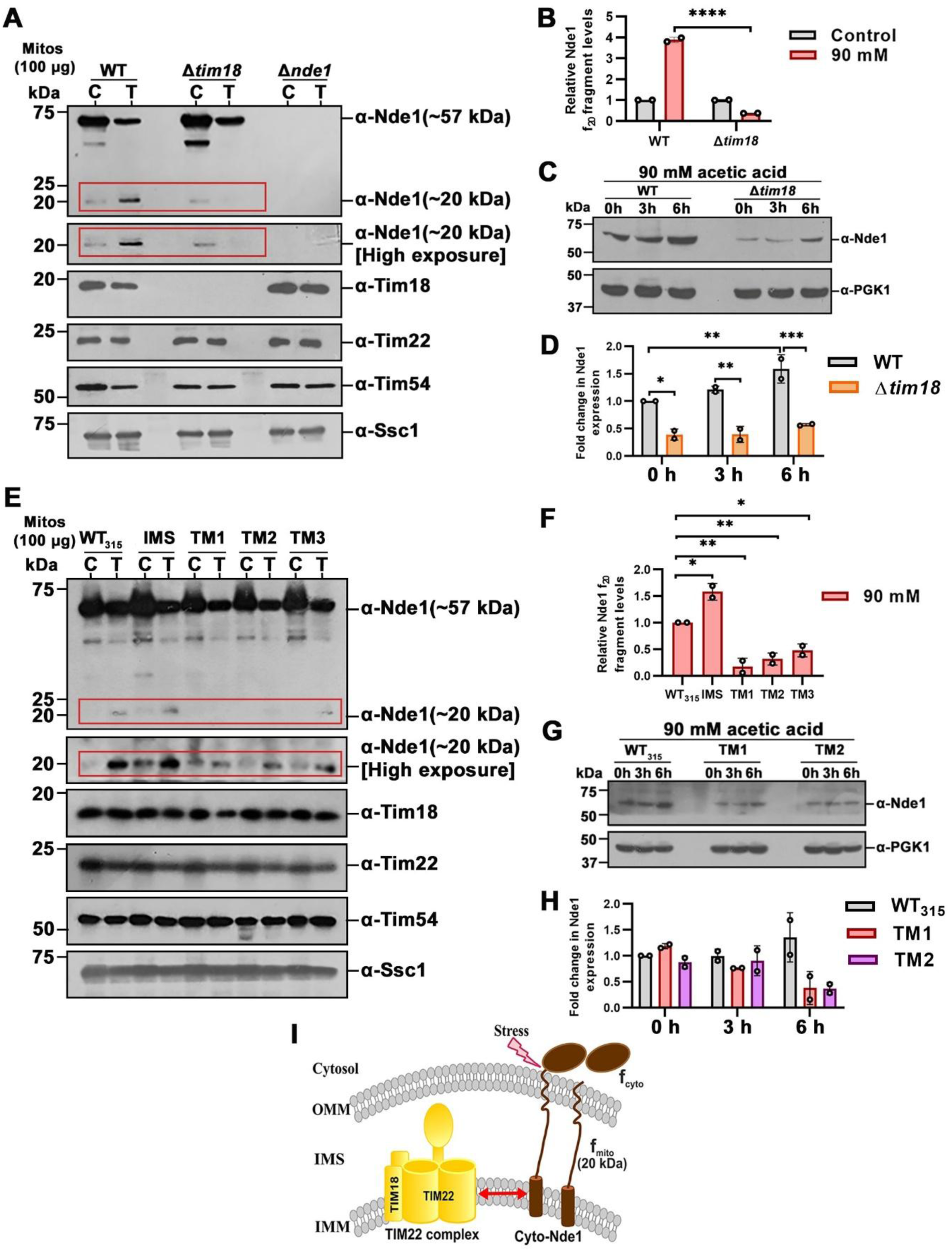
The functional TIM22 complex stabilises the Cyto-Nde1 under stress. (**A, E**) Levels of Nde1 f_mito_ fragment in the deletion and mutant background strains. Mitochondria from untreated and 90 mM acetic acid-treated cells of WT, Δ*tim18,* Δ*nde1* and Δ*tim18*Δ*nde1* and the Tim22 mutants were isolated and analysed using SDS PAGE followed by western blotting using anti-Nde1 and Tim22 complex protein antibodies. Red marked boxes highlight the f_mito_ fragment of Cyto-Nde1. (**B, F**) Levels of the f_mito_ fragment of Nde1 were quantified and represented as fold changes with respect to WT and WT_315_ as the control. Values of *p* ≤ 0.05 were considered significant (*), *p* ≤ 0.01 very significant (**), *p* ≤ 0.001 extremely significant (***) and *p* ≤ 0.0001 very extremely significant (****). (**C, G**) Cytosolic Nde1 levels under stress were analysed in strains with impaired Tim22 complex compared with the WT strains. Cytosolic fractions devoid of mitochondria were isolated from the indicated strains and analysed by immunoblotting (**D, H**). Nde1 protein levels were quantified using Multi Gauge software (n=2). Statistical analysis was performed using two-way ANOVA (with multiple comparisons). Values of *p* ≤ 0.05 were considered significant (*), *p* ≤ 0.01 very significant (**), *p* ≤ 0.001 extremely significant (***) and *p* ≤ 0.0001 very extremely significant (****), with WT at 0h set as the reference. PGK1 was used as a loading control. (**I**) Schematic illustration depicting the Tim22 complex stabilising the Cyto-Nde1 isoform under acetic acid stress.

In correlation with the mitochondrial levels of Nde1 f_mito_ fragment , we next assessed the levels of cytosolic Nde1 fractions at various time intervals upon acetic acid treatment. We collected soluble supernatant fractions devoid of mitochondria (see Method Details) at different time points from WT, Δ*tim18*, TM1, and TM2 strains treated with acetic acid. In wild-type strains (WT and WT_315_) under stress, cytosolic Nde1 levels increased over time, while mutant strains exhibited a significant time-dependent decrease in the cytosolic levels of Nde1, paralleling a reduction in f_mito_ levels (Figs. **5C, 5D** and **5G, 5H**).

### Active TIM22 complex associates with the f_mito_ fragment of Cyto-Nde1

Recognising the significance of Cyto-Nde1 formation under apoptotic conditions, we sought to ascertain if the physical association between the TIM22 complex and Nde1 is a prerequisite. If so, what molecular mechanism governs this interaction during apoptotic processes? This prompted us to perform a co-immunoprecipitation experiment to assess the physical interaction between Nde1 and the TIM22 complex, particularly under stress conditions. Mitochondrial lysates from all the mentioned untagged and Tim18-FLAG tagged strains were subjected to immunoprecipitation under acetic acid-treated and untreated conditions. Intriguingly, both the mature form of Nde1 (∼57 kDa) and cleaved f_mito_ fragment (∼20 kDa) of Cyto-Nde1 were detected in the pulldown fractions (Figs. **6A** and **6C**). However, quantification of the f_mito_ fragment levels revealed a prominent enrichment of this fragment in all the treated samples, except in the TM1 and TM2 mutant strains, thereby highlighting the crucial role of these transmembrane domains in Cyto-Nde1 stabilisation (Fig. **6D**). The presence of IMS-Nde1 in the pulldown fraction suggested a dynamic interaction with the functional TIM22 complex, similar to that observed in Fig. **3E**. Interestingly, under acetic acid treatment, the f_mito_ fragment of Cyto-Nde1 formed a stable, stress-dependent association with the TIM22 complex, suggesting that this processed mitochondrial fragment acts as the key interacting partner of Cyto-Nde1 during apoptosis (Figs. **6A** and **6C**). Taken together, our data indicate that the interaction between the TIM22 complex and the f_mito_ fragment of Nde1 is indispensable for facilitating apoptosis (Fig. **6B**).

**Figure 6.**
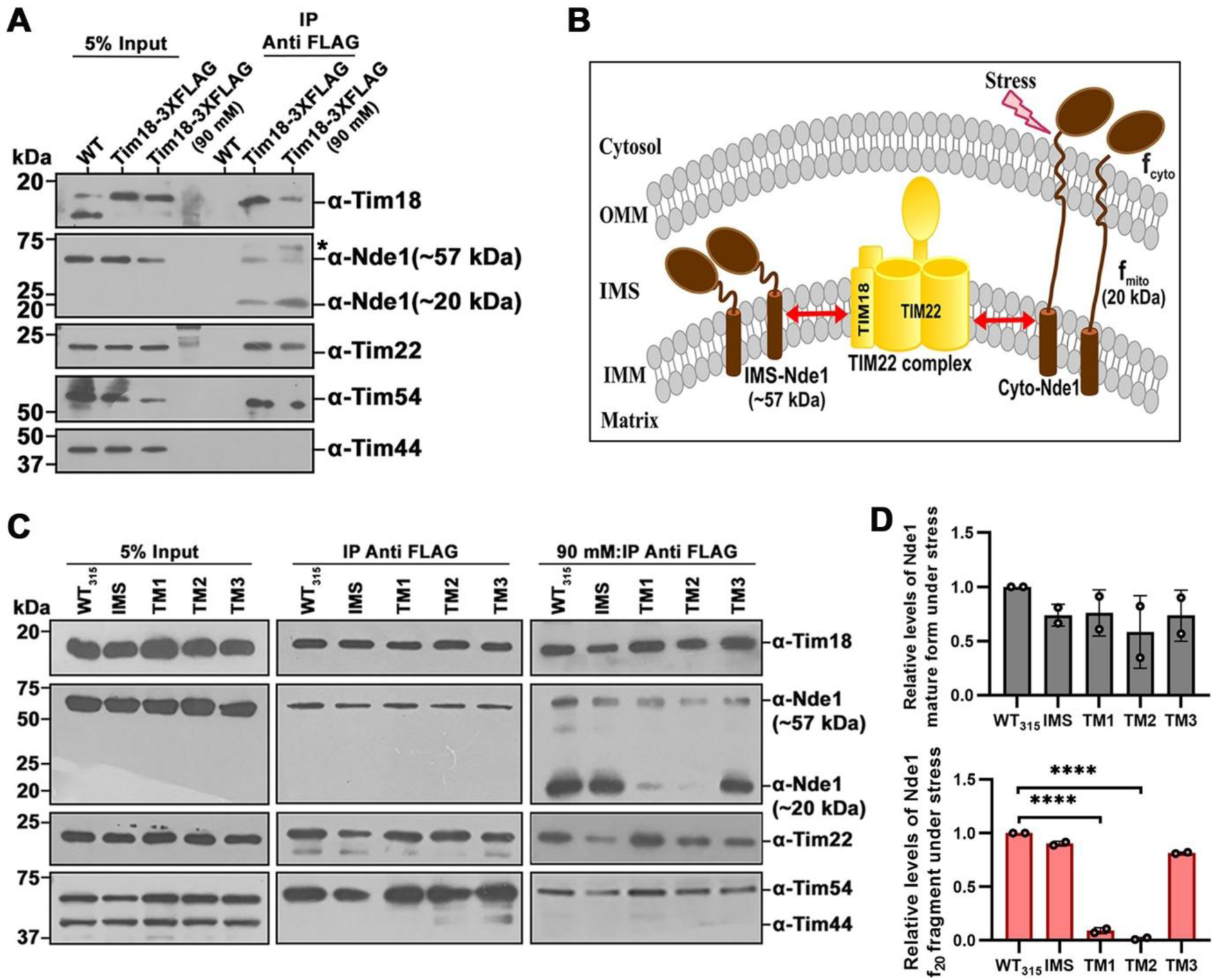
Nde1 regulates TIM22 complex cell death activity via protein-protein interaction. (**A**) Interaction analysis between Tim18 and Nde1 by Co-IP assay. Mitochondria isolated from untagged and Tim18 3X FLAG-tagged WT strains under control and 90 mM acetic acid were solubilised in 1% digitonin buffer and incubated with anti-FLAG conjugated protein G beads. The overnight-bound lysate was then analysed by immunoblotting with antibodies specific to Nde1 and TIM22 complex proteins. 5% of the mitochondrial lysate was loaded as a control. Asterisk indicates the position of IgG heavy chain. (**B**) Schematic representation of the TIM22 complex with both the isoforms of the Nde1 protein. (**C**) Immuno-pulldown assay of the Tim22 mutants under 90 mM stress was performed by solubilising mitochondria from both untagged and 3X FLAG-tagged strains in 1% digitonin buffer, followed by incubation with anti-FLAG conjugated Protein G Sepharose beads. Bound proteins were subsequently analysed by immunoblotting. (**D**) Levels of the Nde1 mature form (IMS-Nde1 of ∼57 kDa) and f_mito_ fragment (∼57 kDa) obtained from panel **6C** were quantified and represented as the fold change concerning WT_315_.

### The combined impairment in the TIM22 complex and Nde1 restores mitochondrial functionality under stress

Perturbations in mitochondrial proteins alter mitochondrial function and organelle dynamics (San-Millán, 2023). In our study, this alteration was induced by acetic acid to examine the effects of the TIM22 complex and Nde1-mediated cell death on mitochondrial parameters. Budding yeast cells, when exposed to acetic acid, either under environmental or industrial conditions, exhibit features resembling intrinsic apoptosis, including the generation of reactive oxygen species (ROS) (Chaves *et al*, 2021; Falcone & Mazzoni, 2016; Giannattasio *et al*, 2008). Thus, to evaluate the changes in mitochondrial morphology under acetic acid stress, mitochondria in the deletion and mutant strains were visualised using a fluorescence microscope with an MTS-mCherry containing plasmid. In untreated conditions, all the cells exhibited reticular and intermediate mitochondrial morphology (Figs. **EV5A** and **EV5B**). However, upon treatment with acetic acid, WT and WT_315_ strains displayed fragmented mitochondria. However, deletion of Tim18 or Nde1 partially restored the morphology. Interestingly, the double deletion strain and the TM1 and TM2 mutant strains completely restored mitochondrial morphology under stress conditions (Figs. **7A** and **7B**).

**Figure 7.**
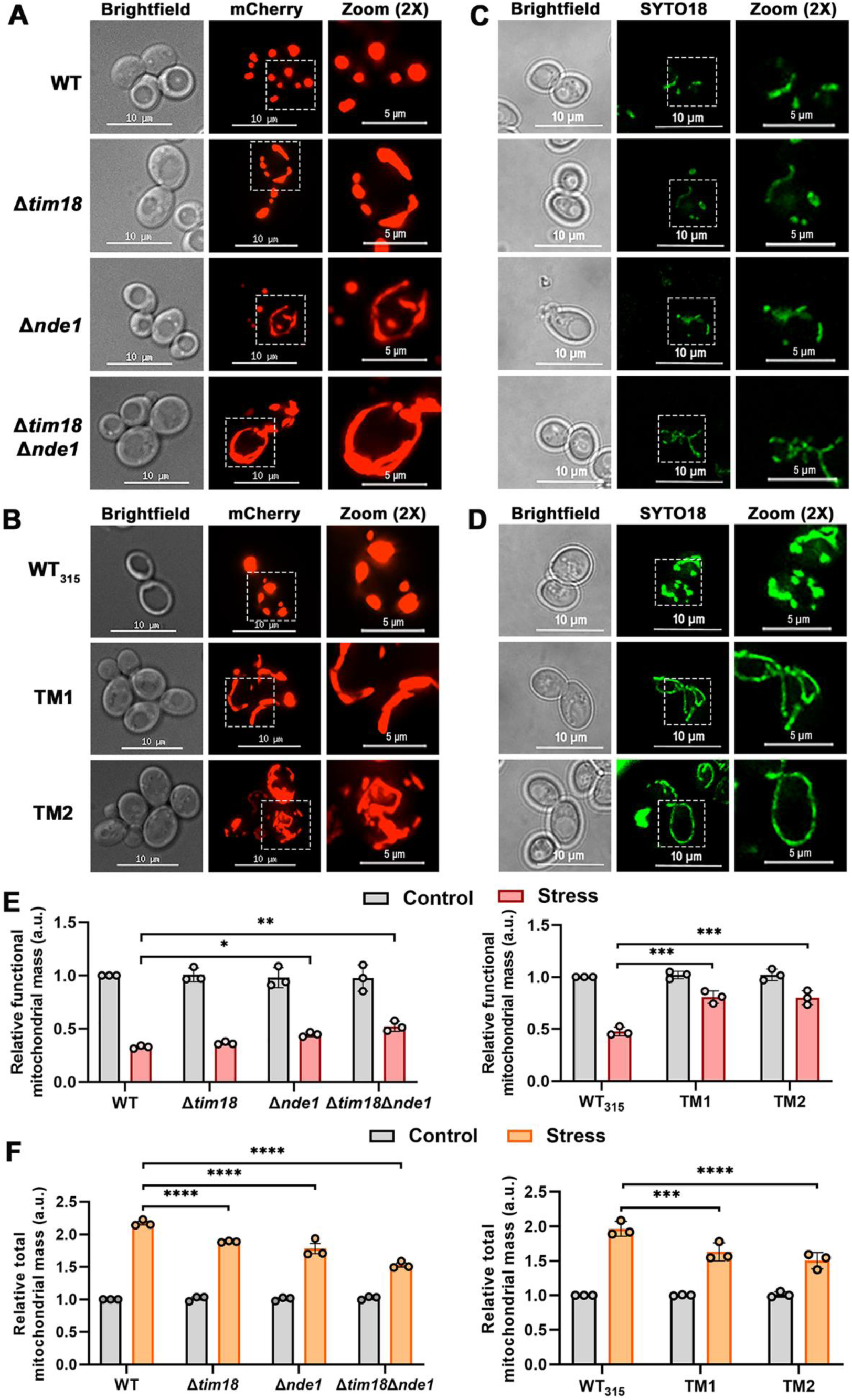
Disruptions in the TIM22 complex and Δ*nde1* cells restore mitochondrial health parameters under acetic acid stress. (**A, B**) Representative images of mitochondrial morphology in the deletion and mutant strains expressing MTS-mCherry marker protein. Cells were grown until mid-log phase in SC-Dextrose media, treated with acetic acid, then immobilised on 2% agarose pads. Images were acquired using a DeltaVision Elite fluorescence microscope equipped with a 100X objective, which was subsequently deconvoluted and processed using SoftWoRx 6.1.3 software. (**C, D**) Representative images of mitochondrial DNA (mtDNA) stained with SYTO18 in the above-mentioned strains. The cells at mid-log phase in SC-Dextrose media were stained with 10 µM for 15 min, and mtDNA was visualised using a Floview 3000 Olympus confocal microscope and processed in ImageJ software. The scale bar represents 10 µm. (**E, F**) Flow cytometry was used to quantify the total and functional mitochondrial mass in the deletion and mutant strains under control and stress conditions, and the results were represented as bar graphs. The strains were stained with 10 μM NAO for 20 min or 8.75 μM TMRE for 15 min to measure total and functional mass, respectively, followed by analysis using BD FACSVerse flow cytometer. The data was analysed using FlowJo software. Quantification data were obtained from three independent experiments (n = 3). In each experiment, 30000 cells were analysed. Values of *p* ≤ 0.05 were considered significant (*), *p* ≤ 0.01 very significant (**), *p* ≤ 0.001 extremely significant (***) and *p* ≤ 0.0001 very extremely significant (****).

Earlier studies have also reported a loss of mitochondrial DNA (mtDNA) stability upon impairment of the TIM22 complex (Kumar *et al*., 2023). Thus, we visualised mtDNA in all the strains using SYTO18 dye. The cells showed intact mtDNA in control conditions (Figs. **EV5C** and **EV5D**). WT cells treated with ETBr (*ρ^-^*) were used as a negative control (Fig. **EV5E**). However, both WT and WT_315_ exhibited fragmented and diminished mtDNA under stress, in agreement with prior studies (Susarla *et al*, 2023). Conversely, the single- and double-deletion strains of Tim18 and Nde1, as well as the TM1 and TM2 mutants, displayed improved mtDNA stability (Figs. **7C** and **7D**).

In addition, we assessed functional and total mitochondrial mass in deletion and mutant strains using TMRE and NAO, respectively, and analysed them by flow cytometry. WT and WT_315_ cells showed reduced mitochondrial membrane potential compared to the single and double deletion strains, as well as the TM1 and TM2 mutants (Fig. **7E**). Surprisingly, an increase in total mitochondrial mass was observed in all the strains under acetic acid stress, despite the fragmented mitochondrial morphology and reduced mitochondrial potential. However, the increase was significantly reduced in the double deletion and TM1, TM2 mutants compared to their respective wild-type strains (Fig. **7F**). This increase in total mass could be a compensatory response initiated by mitochondria under oxidative stress conditions, in which mitochondria increase their overall mass while maintaining a fragmented morphology (Oliveira *et al*, 2015).

Defective import complex and loss of mitochondrial potential are often associated with increased ROS generation in mitochondria (Zorov *et al*, 2014). Thus, ROS generation in all the studied strains was assessed using MitoSOX staining. Under stress, the Δ*tim18*Δ*nde1* as well as the TM1 and TM2 mutant strains exhibited a lesser number of cells with MitoSOX staining as compared to the WT and WT_315_ cells (Figs. **8A** and **8D, 8B** and **8E**). The relative median intensity, as assessed by flow cytometry, also yielded similar results (Figs. **8C** and **8F)**. All these findings suggest that the functional TIM22 complex, mainly the Tim18 subunit along with the TM1 and TM2 domains, plays a vital role in mediating apoptosis under stress and in protecting overall mitochondrial health.

**Figure 8.**
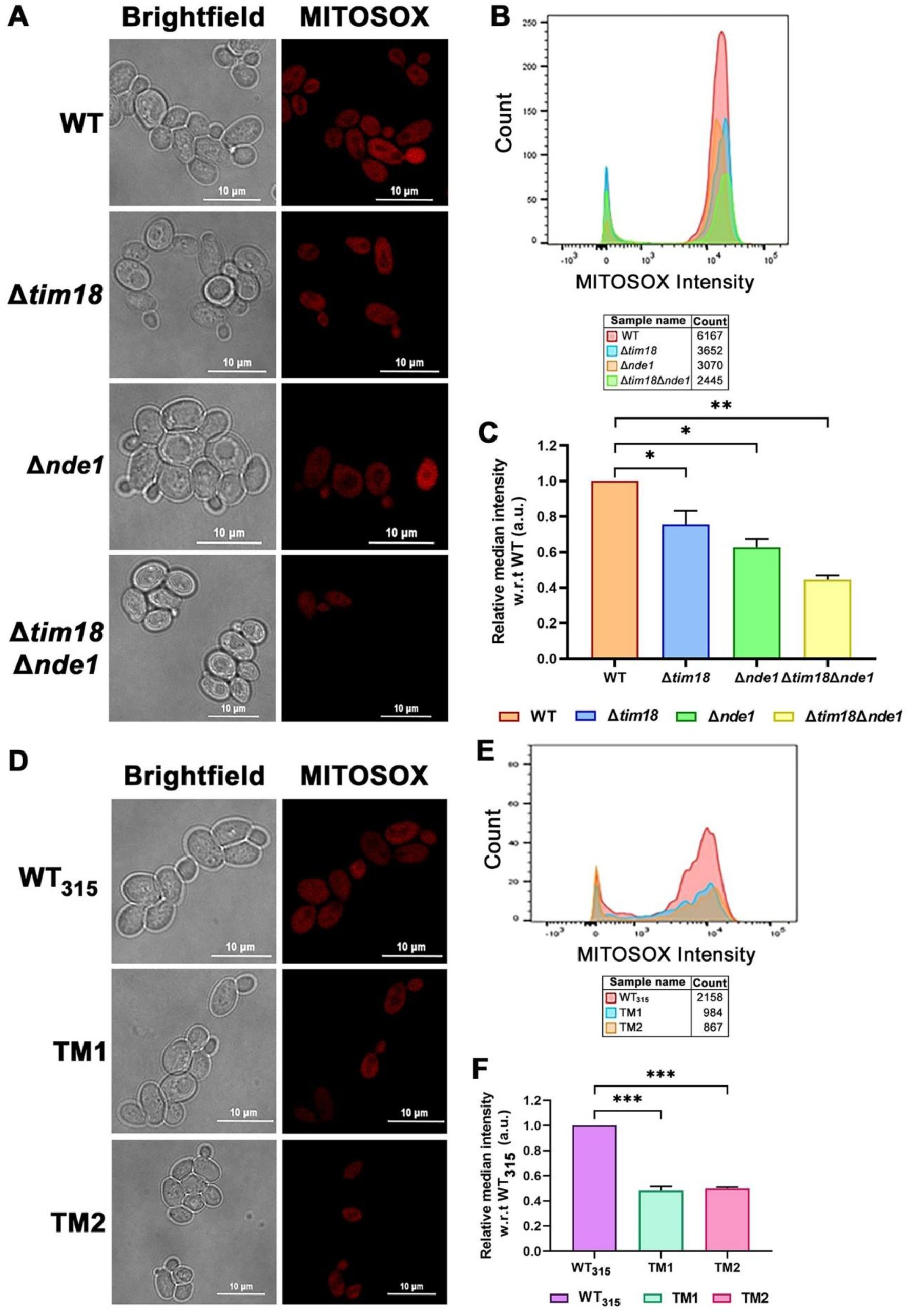
Mutations in the TIM22 complex and the absence of Nde1 reduce mitochondrial ROS under acetic acid stress. (**A, D**) Representative microscopy images depicting mitochondrial ROS levels in cells deleted for Nde1 and with mutations in the Tim22 subunits stained with MitoSOX dye, respectively. The cells grown to mid-log phase in SC-Dextrose media at 30°C were treated with acetic acid and subsequently stained with 10 µM MitoSOX. Images were acquired using an Olympus Fluoview 3000 confocal microscope. Scale bar represents 10 µm. (**B, E**) MitoSOX fluorescence intensities were quantified in 30 randomly selected cells using ImageJ and represented as line profiles. (**C, F**) Flow cytometry analysis of mitochondrial ROS accumulation using the MitoSOX dye was performed on 30,000 cells per sample from two independent experiments (n = 2). Statistical analysis was performed using two-way ANOVA (with multiple comparisons) Values of *p* ≤ 0.05 were considered significant (*), *p* ≤ 0.01 very significant (**), *p* ≤ 0.001 extremely significant (***) and *p* ≤ 0.0001very extremely significant (****).

In summary, we propose a model which depicts that under acetic acid stress, the functional TIM22 complex stabilises the Cyto-Nde1 topomer, specifically through its interaction with the f_mito_ fragment. This complex stabilisation facilitates activation of the apoptotic pathway in the cell, leading to mitochondrial DNA fragmentation and increased ROS production. On the contrary, disruption or mutation in the TIM22 complex impairs the stabilisation of Cyto-Nde1, thereby suppressing the apoptotic response and restoring mitochondrial integrity (**Fig. 9**).

**Figure 9.**
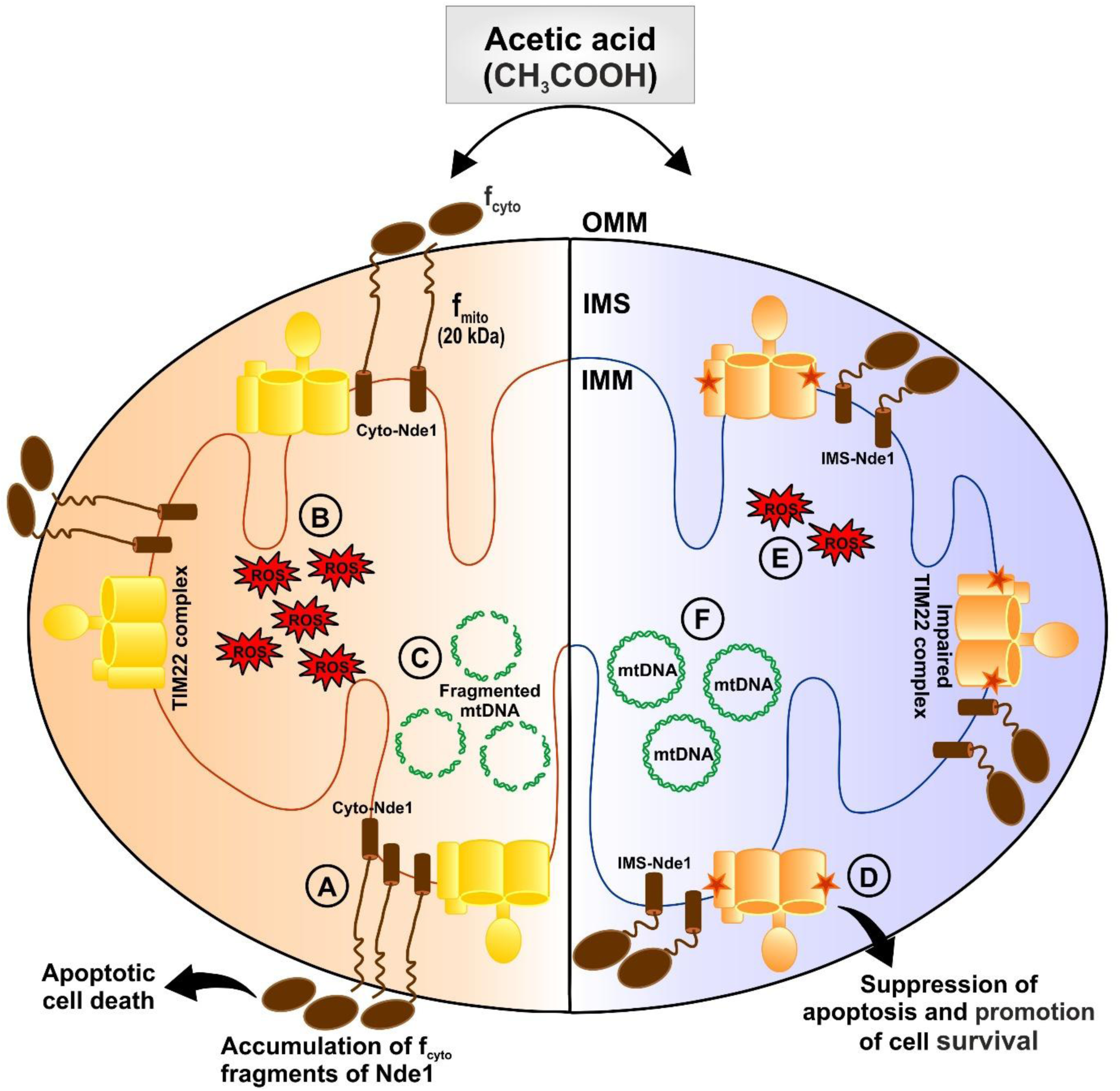
Summary model illustrating the apoptotic role of the TIM22 complex in association with the topological dynamics of Nde1. (**A**) Upon acetic acid stress, the functional TIM22 complex stabilises the Cyto-Nde1 through its f_mito_ (20 kDa) fragment, thereby leading to apoptosis. (**B, C**) An active TIM22 complex facilitates the acetic acid-induced apoptotic signaling, leading to elevated mitochondrial ROS levels and mitochondrial DNA (mtDNA) fragmentation. (**D**) Impairment of the TIM22 complex suppresses apoptotic induction under acetic acid stress, favouring the activity of the IMS-Nde1 isoform. (**E, F**) A dysfunctional TIM22 complex attenuates the stress response, reducing ROS accumulation and preserving mtDNA integrity, thereby restoring overall mitochondrial function.

## Discussion

Programmed cell death, or apoptosis, is an evolutionarily conserved process across eukaryotic lineages, ranging from unicellular to complex multicellular species (Newton *et al*, 2024). In yeast, apoptosis may serve as a survival mechanism which eliminates a significant portion of the population under starvation or stress conditions, thereby favouring the survival of fitter cells (Büttner *et al*, 2006). Multiple factors contributing to apoptosis in yeast have been identified: the precise molecular mechanisms underlying mitochondrial involvement remain largely uncharacterised. External stress stimuli often trigger apoptosis, leading to mitochondrial outer membrane permeabilisation (MOMP) (Dadsena *et al*., 2021). This leads to the release of mitochondrial ETC proteins, such as cytochrome c, into the cytosol (Du *et al*., 2007; Kalpage *et al*, 2020; Wang, 2001). Several proteins of the mitochondrial import complex machinery have recently evolved as crucial secondary mediators of apoptosis. Mitochondrial translocases, such as the Tim50 subunit of TIM23 translocase and Tom20 subunit of TOM complex, potentially interact with pro-apoptotic Bcl-2 proteins, thereby highlighting the moonlighting roles of the import complexes in apoptotic regulation (Guo *et al*, 2004; Lalier *et al*, 2021). A recent study has highlighted the role of Nde1 (a component of the yeast ETC) as a pro-apoptotic factor, primarily affecting mutants of the carrier translocase or the TIM22 complex (Saladi *et al*., 2020). However, the interplay between ETC and the protein import machinery in regulating apoptosis remains elusive.

In the present study, we demonstrate a vital role for the TIM22 complex in cell death, mediated primarily by the Tim18 subunit and the transmembrane regions 1 and 2 of the Tim22 core protein. A mutation in the IMS region of Tim22 destabilises the entire TIM22 complex (thiol-reduced Tim22 protein) and thereby decreases stress tolerance. The reactive thiol (-SH) groups of cysteine residues, which form disulfide bonds (S-S) in proteins, act as redox switches and are thus sensitive to diverse stress stimuli (Okamoto *et al*, 2014). Additionally, a mutation in the conserved residue of the TM3 domain is associated with defects in the import of Tim22 substrates (Kumar *et al*., 2020). Under stress, this impaired translocase activity resulted in a more severe growth phenotype. However, mutations in the TM1 and TM2 domains did not disrupt disulfide bridge stability, thereby suppressing the stress-sensing mechanism. Interestingly, these TM1 and TM2 mutants, as well as the Δ*tim18* strain lacking the *NDE1* gene, further suppressed the stress-induced growth defects (Fig. **1**). This implies a genetic interaction between the carrier translocase and Nde1 that facilitates apoptosis under stress conditions.

Thus, under acetic acid stress, a functional TIM22 complex stabilises the Cyto-Nde1 isoform, which subsequently promotes the accumulation of f_cyto_ fragments of Nde1, ultimately leading to cell death. In contrast, destabilisation of the TIM22 complex, either through deletion or mutation, suppresses this stress-sensing mechanism, potentially leading to cell survival and uncontrolled cell proliferation (Fig. **8****; points A and D**) (Jackson *et al*, 2021). This dual functionality of the TIM22 complex in yeast is analogous to the AGK subunit of the mammalian TIM22 complex, which similarly exhibits bifunctional activity in mitochondrial protein biogenesis and phospholipid homeostasis (Vukotic *et al*, 2017).

Further supporting this model, Nde1 overexpression in the Δ*tim18* strain and in the TM1 and TM2 mutants resisted acetic acid stress and reduced cytotoxicity. The reverse condition did not elicit cellular damage (Fig. **3** and **EV3**). A possible reason is that the TIM22 complex acts upstream of Nde1 in the apoptotic pathway. This parallels the PINK1-Parkin pathway in mitochondrial quality control. In this pathway, PINK1 accumulates on damaged mitochondria and signals for Parkin-mediated degradation (Green *et al*, 2010). Such interactions provide a regulated mechanism for apoptosis, preventing aberrant cell death.

Apoptosis is often characterised by elevated mitochondrial ROS levels and compromised mtDNA integrity, which ultimately disrupts overall mitochondrial homeostasis (Ježek *et al*, 2018). Functional TIM22 complex and the Nde1 protein responded to the acetic acid stress, which upregulated mitochondrial ROS production and mtDNA fragmentation (Fig. **8****; points B and C**). However, an impaired carrier complex exhibited an attenuated stress response, leading to reduced ROS accumulation and preservation of mtDNA integrity, thereby restoring overall mitochondrial function (Fig. **8****; points E and F**).

The association between the TIM22 complex and Nde1 in modulating the cell death pathway under stress conditions provides a clue into the evolutionary conservation of the TIM22 complex’s functional dependency on the respiratory chain in higher eukaryotes. This interaction between the carrier translocase and the ETC serves as a well-orchestrated, highly regulated apoptotic pathway. This raises the possibility that this unconventional cell death pathway involving the Nde1 and TIM22 complex in yeast could also be conserved across phylogeny. The genetic variants in the subunits of the TIM22 complex have been linked to several pathologies, underscoring the importance of this complex in maintaining mitochondrial health and cellular viability. Further research into the role of the TIM22 complex in human apoptosis in mammals could provide valuable insights into the mechanisms of apoptosis and development of potential therapeutic strategies (Mussulini *et al*, 2025; O’Malley *et al*, 2020; Weiss *et al*, 2022).

## Method Details

### Reagents and tools table

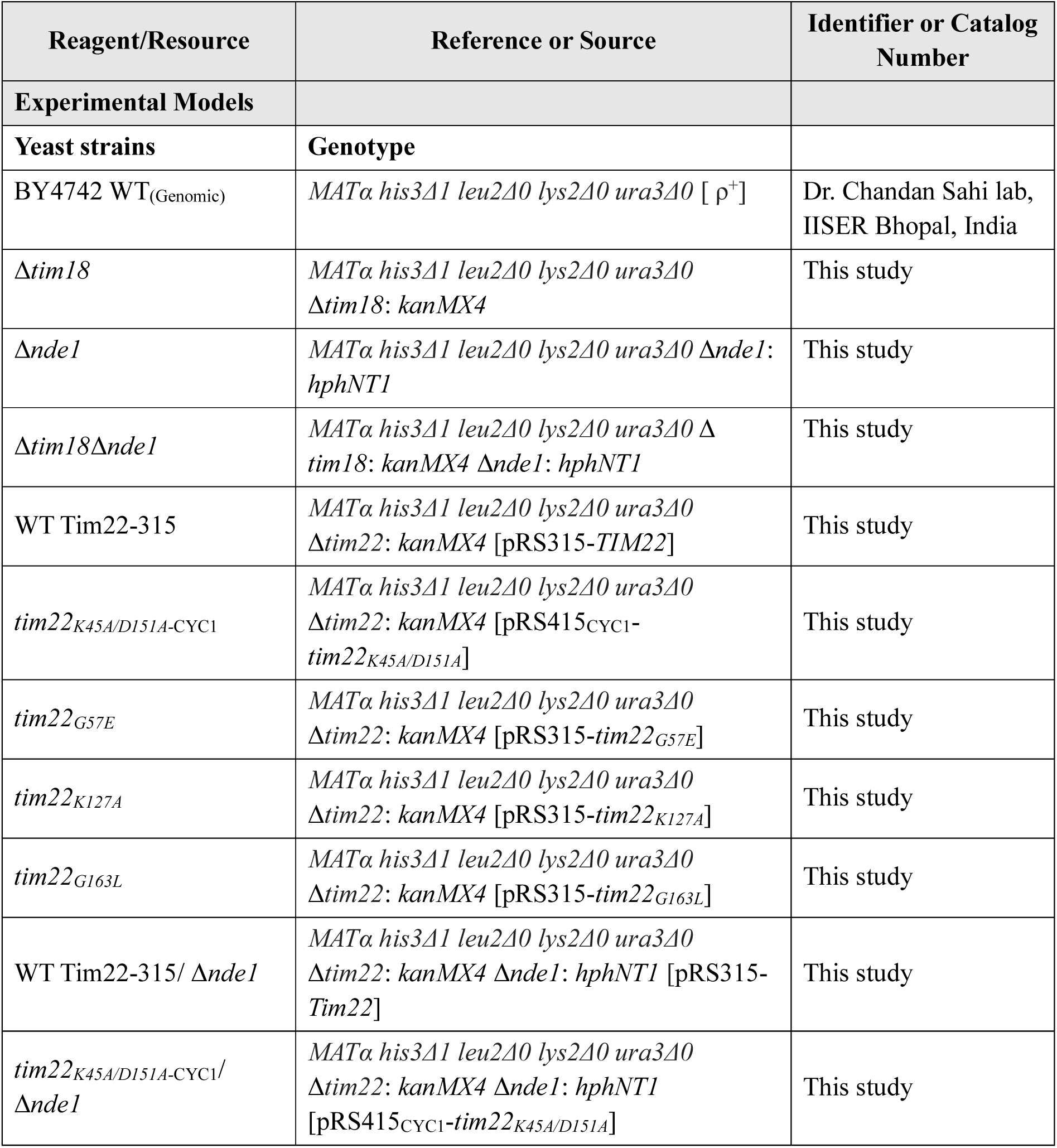
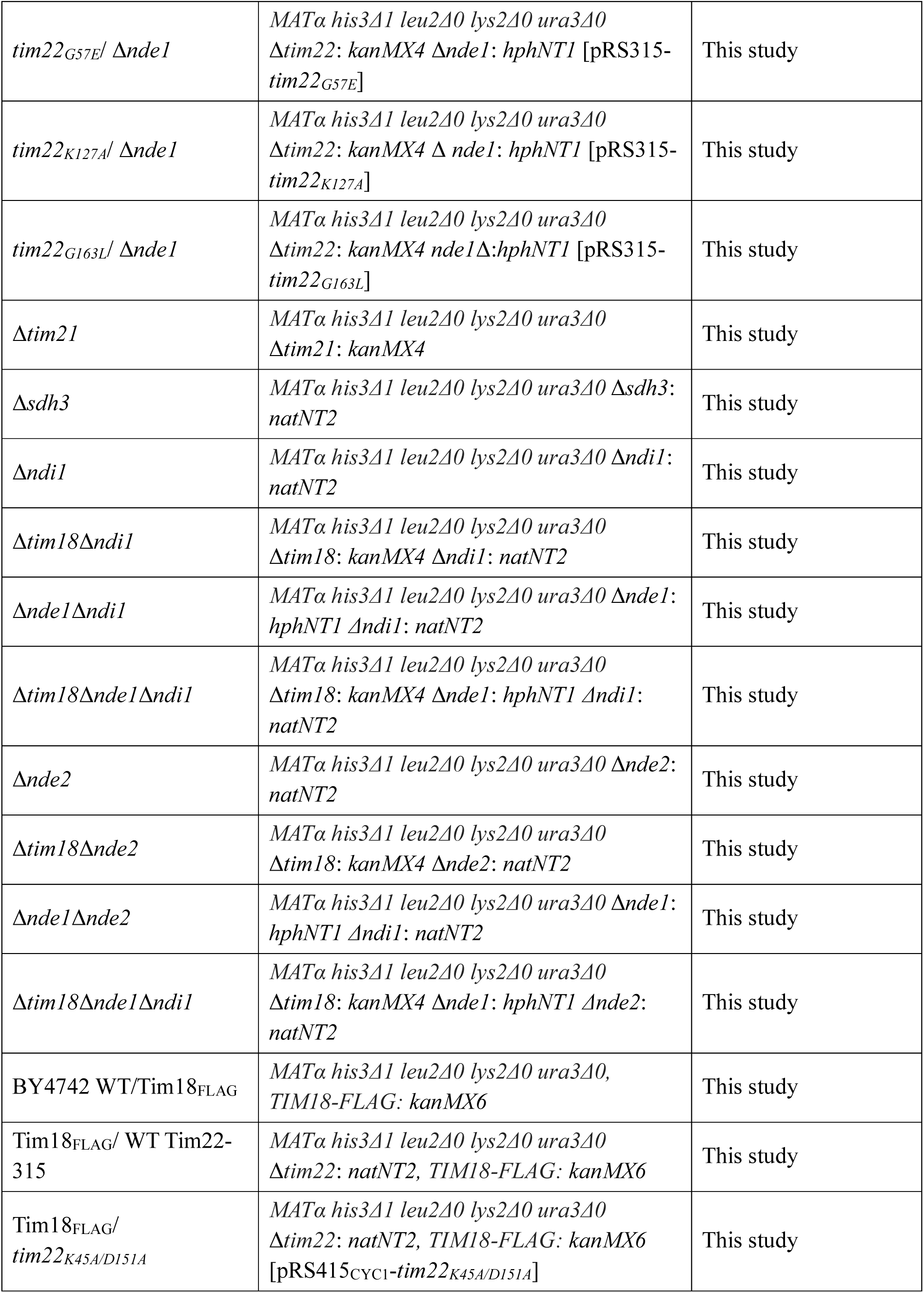
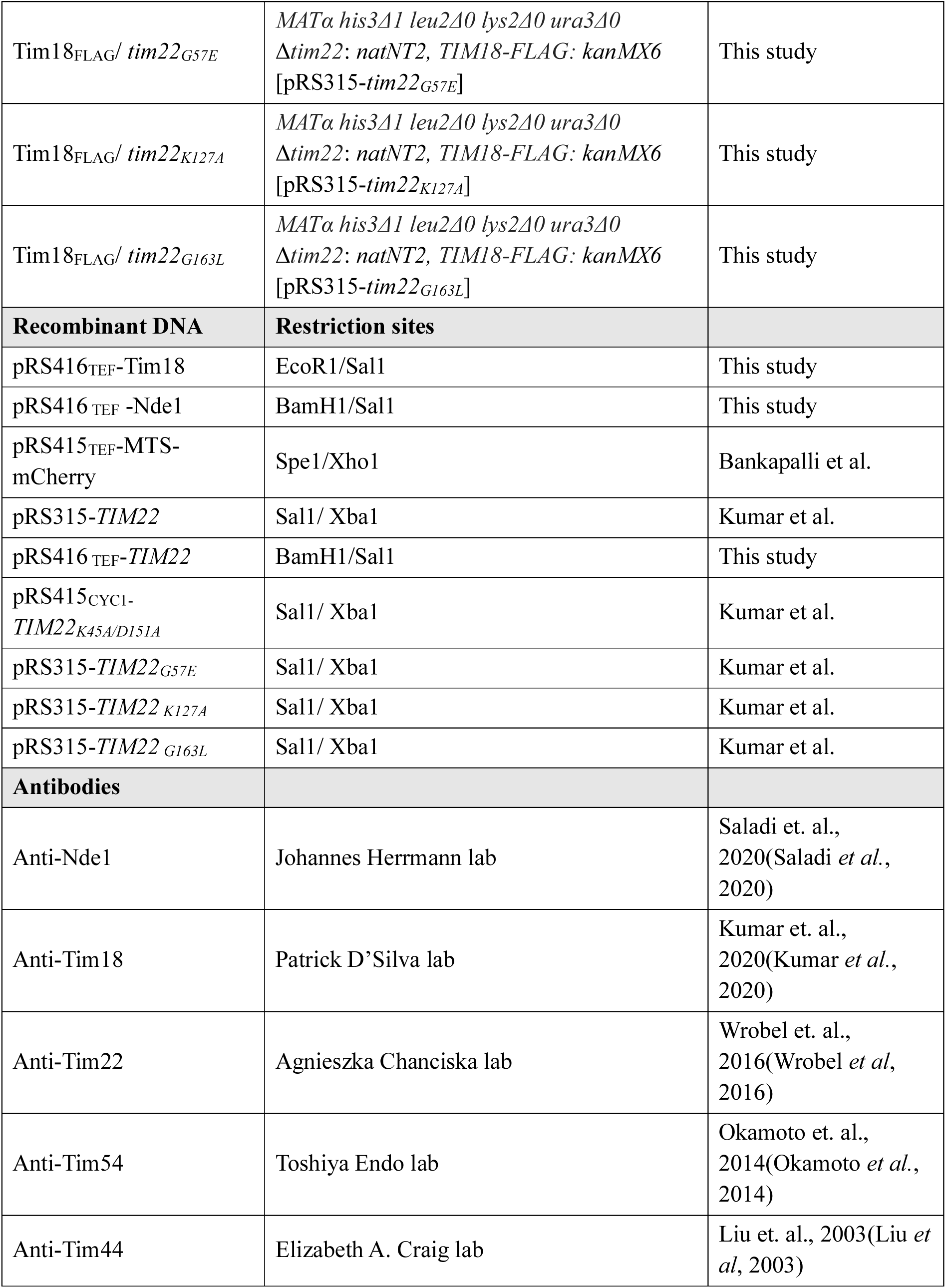
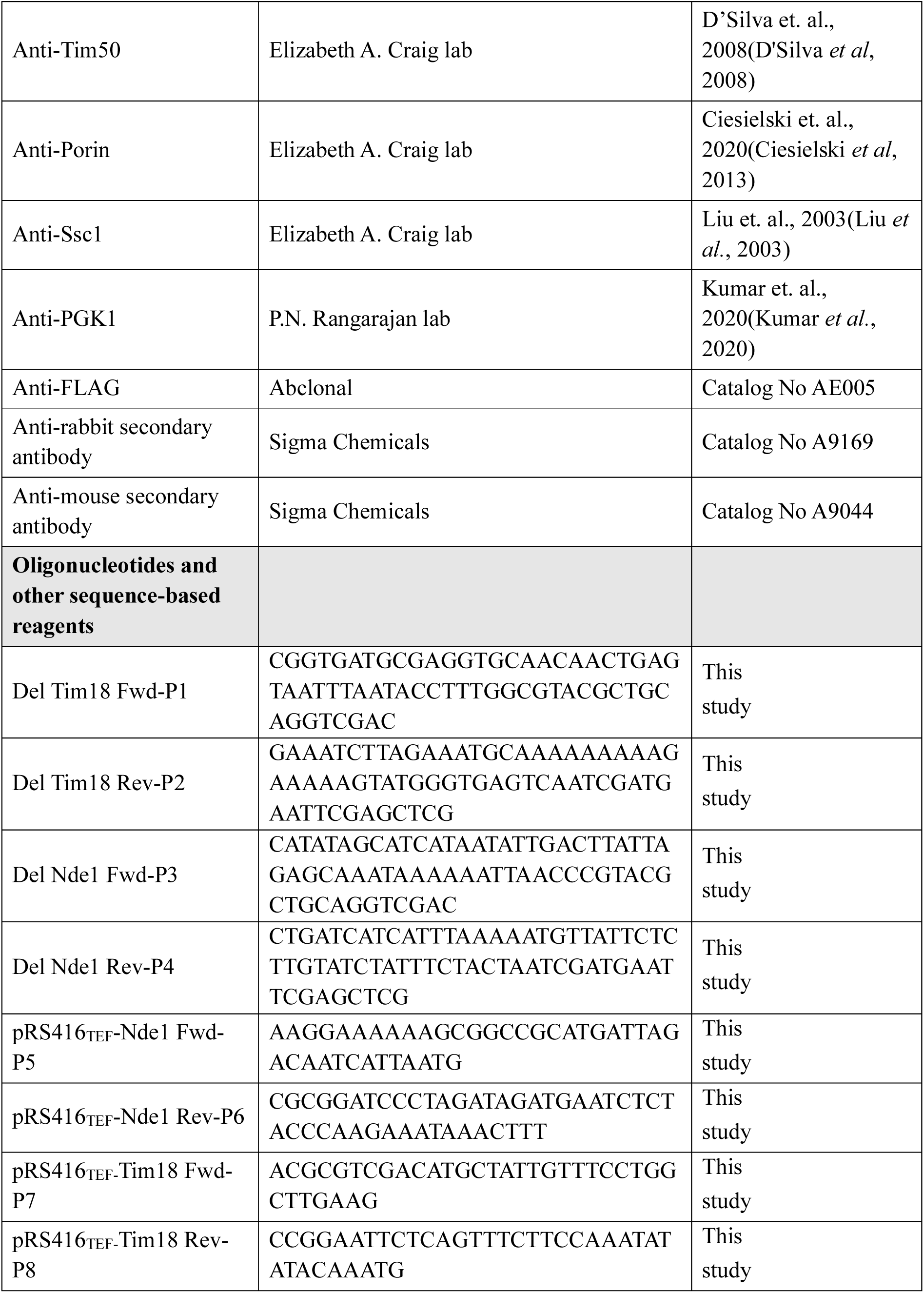
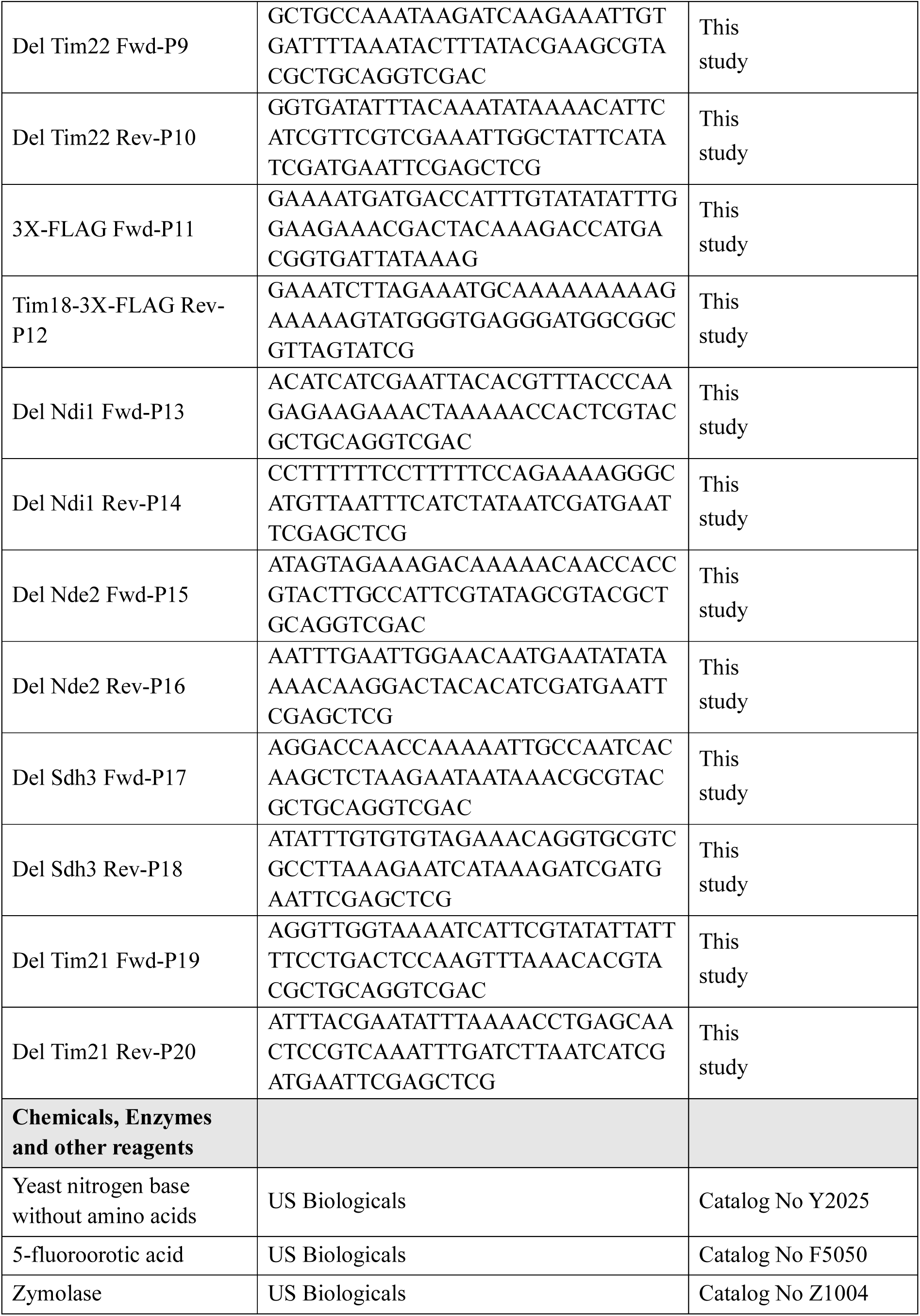
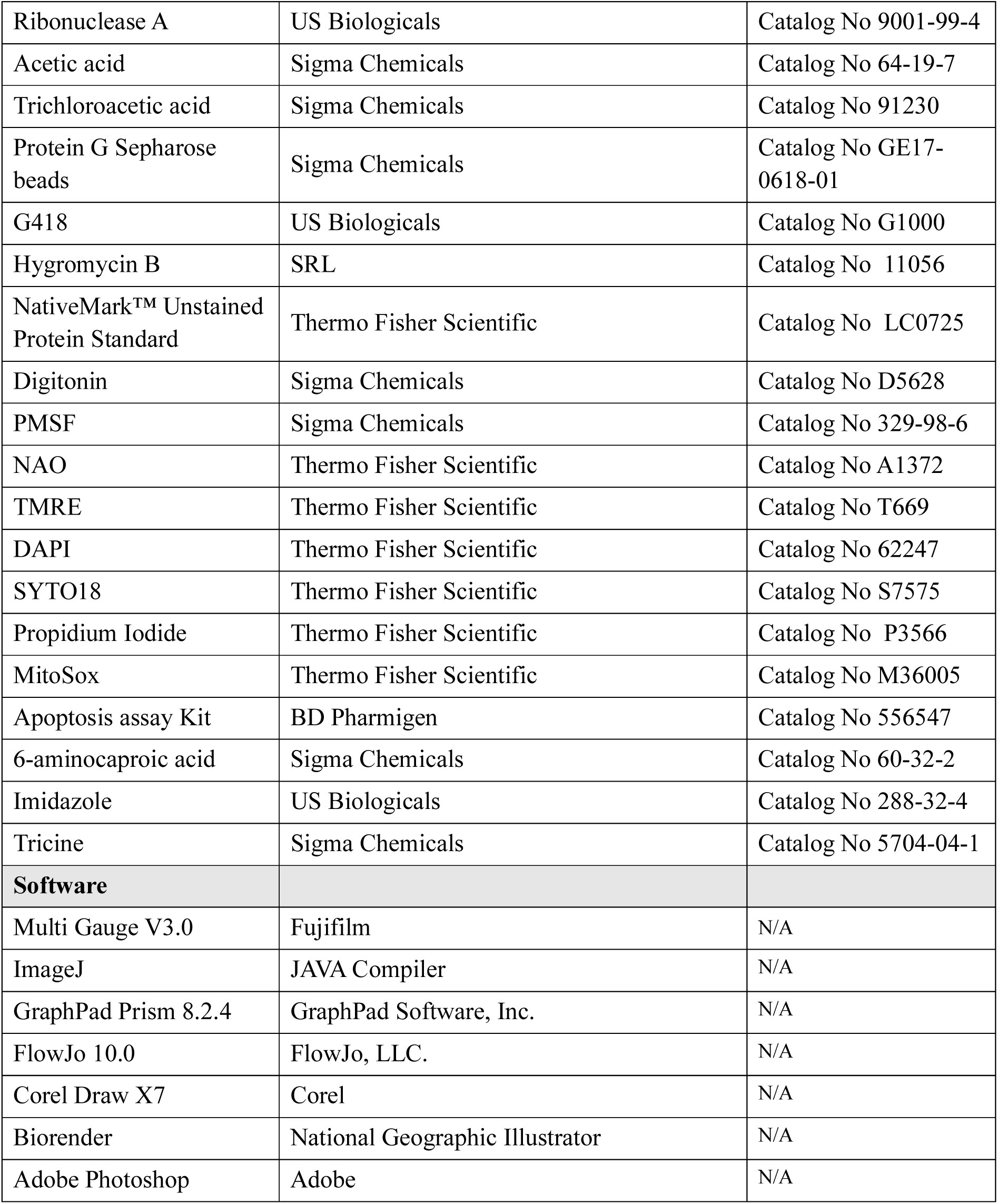

#### Construction of yeast strains and plasmids by genetic technology

*Saccharomyces cerevisiae* strains used in this study are listed in **Table S1**. All primer sequences used for PCR amplification are listed in **Table S2**. Genes cloned under the shuttle vectors are detailed in **Table S3** of the resource detail section.

Yeast strains used in this study are based on the BY4742 haploid Mat-α background. All deletion strains in this study, including Δ*tim22*, Δ*tim18*, Δ*sdh3*, and Δ*tim21*, were generated by PCR-based homologous recombination to substitute the entire ORF with a *KanMX* selection cassette. Whereas Δ*nde1* was replaced with *Hph* (hygromycin) selection marker, Δ*nde2,* Δ*ndi1,* and Δ*tim22* were generated using *natNT*2 cassettes. All primers are listed in **Table S2**.

The PCR amplified product (*TIM22*) was cloned into centromeric vector pRS315, which carries the *LEU2* auxotrophic marker (gift from Prof. Elizabeth A. Craig, University of Wisconsin Madison). Tim22 mutants were generated by random mutagenesis, then cloned into the pRS315 and pRS415_CYC1_ vectors using Xba1/Sal1(Kumar *et al*., 2020). Firstly, pRS416-*TIM22* was transformed into WT using the lithium acetate (LiAc) method. Genomic *TIM22* was deleted using P9 and P10 primers(Daniel Gietz & Woods, 2002). Subsequently, the pRS315-*TIM22* and the mutants encoded with the *LEU2* auxotroph marker were transformed into the WT Tim22-416 strain. Strains expressing only *TIM22* encoded by pRS315 and pRS415 were selected by patching transformants onto 5-fluroroorotic acid (5-FOA) plates. The strains were streaked on leucine and uracil dropout media to check the persistence of the *LEU2* marker while the *URA3* marker was lost.

The *TIM18* and *NDE1* genes were amplified from genomic DNA using the forward primers P5 and P6, and the reverse primers P7 and P8, respectively. After amplification, the TIM18 fragment was cloned into the EcoR1/Sal1 digested yeast vector pRS416_TEF_. The Nde1 fragment was cloned into the BamH1/Sal1 digested yeast vector pRS416_TEF_, which contains *URA3* as an auxotrophic marker.

The TIM18 gene was chromosomally tagged with a FLAG epitope at the C terminus by homologous recombination for expression analysis. The DNA fragment encoding 3X-Flag and flanking regions homologous to recombination with the TIM18 loci was PCR-amplified using ED-15 as a template and primers P11 and P12. Positive transformants were selected on a media containing G418 (*KanMX*4 cassette- 400 μg/μl).

### Growth media and stress conditions

*S. cerevisiae* cells were either grown in (i) YPD media containing yeast extract (1%), peptone (2%), dextrose (2%), (ii) Complete synthetic minimal media or CSM media containing yeast nitrogen base without amino acids (0.67%), complete synthetic media (0.072%) and dextrose (2%) or (iii) synthetic complete dropout (leucine, uracil or both leucine and uracil) media *i.e.* SCD-Leu, SCD-Ura or SCD-Ura-Leu media containing yeast nitrogen base without amino acids (0.67%), leucine dropout supplement (0.069%), uracil dropout supplement (0.069%) or double dropout supplement (0.072%) and dextrose (2%). For growth on plates, 3% agar was added to the media. Cells were grown overnight at 30°C and then shifted to fresh media. Cells were allowed to reach the exponential growth phase for all experiments.

For acetic acid stress, the cells were resuspended in sterile water and treated with 50 mM, 70 mM, or 90 mM acetic acid, followed by incubation for 1h or 3h at 30°C. Subsequently, the cells were washed in sterile water and used for further experiments.

### Yeast viability assay (Spot test) and growth curve

The respective yeast strains were grown overnight in liquid YPD, synthetic complete or dropout media for spot analysis. Total yeast cells, equivalent to an OD_600_ ∼0.5, were harvested at mid-log phase. The cells were resuspended in sterile water and treated with various concentrations of acetic acid (Sigma Chemicals), followed by incubation at 30°C for 3h. Subsequently, the cells were washed in sterile water and subjected to 10-fold serial dilutions. Each dilution was spotted on the respective media plates and incubated at 30°C and 37°C for 36-180h. To obtain growth curves, the cells were grown overnight and diluted to an OD_600_ ∼0.1 in a transparent 96-well plate. Growth kinetics of the untreated and acetic acid-treated samples were monitored at 30°C. 90 mM acetic acid was added to the cultures when the cells reached the mid-log phase, followed by further incubation at 30°C for 48h. By measuring the absorbance at 600 nm at a regular interval of 2h using a Tecan microplate reader and iControl software, the data were fitted with GraphPad Prism 8.2.4 software.

### Apoptotic assay by FITC Annexin V and PI staining

Apoptosis was determined using the FITC AnnexinV apoptosis detection kit (BD Pharmingen), following the protocol described previously(Bankapalli *et al*., 2020). Both untreated and acetic acid-treated cells were washed in sorbitol buffer (1.2 M sorbitol, 35 mM potassium phosphate, 0.5 mM MgCl_2_, pH 6.8) and digested with 15 U/ml zymolase (US Biologicals) at 28°C for 1h. Yeast cells were then harvested, washed with annexin binding buffer and resuspended in the same buffer with 2 μl of FITC Annexin V and 2 μl of propidium iodide (PI). Samples were incubated at room temperature for 20 min before being subjected to flow cytometry analysis.

### Chromatin condensation visualisation by DAPI staining

To assess chromatin condensation, the nuclei of the mentioned strains were stained with DAPI (4,6-diamidino-2-phenylindole)(Li *et al*., 2006). Yeast cells collected at the exponential phase (OD_600_ ∼0.3-0.4) were exposed to 90 mM acetic acid for 1h, followed by brief fixation with 70% ethanol and stained with DAPI (2 µg/ml) for 15 min. Cell images were then acquired with a confocal microscope (Olympus Fluoview 3000). Images were processed and assembled using Adobe Photoshop CC.

### Mitochondrial lysate preparation

Yeast mitochondria were isolated following the previous protocols with slight modifications(Matta *et al*, 2017; Meisinger *et al*, 2006; Pareek *et al*, 2013). Yeast strains cultured in complete synthetic media with and without acetic acid stress were incubated at 30°C. Cells were harvested at 6000 rpm, weighed, and resuspended to 0.5 g wet weight/ml of buffer (100 mM Tris-H_2_SO_4_ pH 9.4, 10 mM dithiothreitol). The suspension was incubated at 25°C for 20 min with gentle shaking (70 rpm), after which the cells were harvested and washed with 1.2 M sorbitol. Subsequently, they were resuspended in zymolase buffer (1.2 M Sorbitol, 20 mM KH_2_PO_4,_ pH 7.4) to obtain a 0.15 g wet weight/ml suspension in buffer. Zymolase was added to the suspension at a concentration of 2 mg/gm of wet weight, and the suspension was incubated at 25°C for 1h with gentle shaking. Spheroplast formation was confirmed by checking cell morphology in 0.1% SDS and 1.2 M sorbitol, and the spheroplasts were then harvested at 5000 rpm, followed by washing with 1.2 M sorbitol once. For homogenisation, the spheroplasts were gently resuspended in a chilled homogenisation buffer (0.6 M sorbitol, 10 mM Tris-Cl, pH 7.4, 0.2% BSA and 1 mM PMSF) to 0.5 g wet weight/ml buffer. Under ice-cold conditions, the suspension was subjected to 30+25 strokes in a douncer with homogenisation buffer, followed by centrifugation at 3000 rpm. The supernatant was stored at 4°C, while the resultant pellet was resuspended in the same volume of homogenisation buffer as before and subjected to another round of homogenisation. The supernatant obtained after centrifugation at 3000 rpm was pooled with the previous supernatant and subjected to centrifugation at 15,000 rpm in an Oakridge tube to obtain a crude mitochondrial pellet. The pellet was resuspended in 1 ml of homogenisation buffer and subjected to cleaning by alternating between 5000 rpm and 13,000 rpm at each resuspension step. Finally, the purified mitochondria pellet was resuspended in 0.75-1 ml SEM buffer (10 mM MOPS KOH, pH 7.2, 250 mM Sucrose, 1 mM EDTA), frozen in liquid nitrogen, and stored at −80 °C until further use.

During the alternating centrifugation step, the supernatant obtained after centrifuging the sample at 13,000 rpm is considered the fraction without mitochondria and is referred to as the cytosolic fraction. This fraction was obtained from wild-type and mutant strains at 0h, 3h, and 6h, and western blot analysis was performed using the respective antibodies.

### Whole-cell lysate extraction

For whole-cell lysate preparation, equal cells of OD_600_ ∼0.4 were harvested and resuspended in the required amount of 10% trichloroacetic acid (TCA). Cells were then incubated on ice for 30 min with intermittent mixing. The pellet obtained after centrifugation was washed twice with ice-cold acetone. To remove excess acetone, the pellet was air-dried at 42°C for 15 min. The dried pellet was resuspended in 1X SDS dye and lysed with acid-treated glass beads. This was followed by vortexing the sample for 10 min and boiling at 90°C for 10 min. The supernatant obtained after centrifugation was loaded on an SDS gel.

### Western blot analysis

The SDS gel was electroblotted into the PVDF membrane in 1X Tris-Glycine-Methanol buffer. Following the transfer, the blot was blocked with 5% skimmed milk for 1h and washed with 1X TBST buffer (Tris base, NaCl, 0.1% Tween-20). The blot was then incubated with primary antibodies, followed by four washes with 1X TBST buffer. Next, the blot was incubated in HRP-linked anti-rabbit secondary antibody (1:15,000/1:10,000 dilution) for 1h and again washed with buffer three times. Luminol reagent mix was added to the blot and exposed to X-ray film.

### Blue Native PAGE (BN-PAGE) Analysis

Blue Native Polyacrylamide gel electrophoresis (BN-PAGE) was performed according to a previously described protocol(Wittig *et al*, 2006). 400-600 μg/ml mitochondria were solubilised in 100 µl of digitonin buffer (1% digitonin, 50 mM NaCl, 50 mM Imidazole, 2 mM 6-aminohexanoic acid, 1 mM EDTA, pH 7.0) for 45 min at 4°C. The solubilised material was centrifuged (14,000 rpm for 15 min at 4°C) to separate soluble and insoluble components, and 15 µl of sample buffer (2.5 µl of 5% Coomassie Brilliant Blue-G and 12.5 µl of 50% glycerol) was added to 100 µl of the supernatant. Samples were loaded onto a 6-16% gradient native imidazole PAGE gel for immunoblotting and subsequently detected using Nde1, Tim54, and Porin-specific antibodies.

### Co-immunoprecipitation studies

Mitochondria (5 mg/ml) were solubilised in 1ml of digitonin buffer (1% digitonin, 25 mM Tris HCl, pH 7.5, 50 mM KCl, 5 mM EDTA, 10% Glycerol and 1 mM PMSF). The digitonin-solubilised fraction was separated by centrifugation and diluted with 500 μl of buffer (25 mM Tris-HCl, pH 7.5, 50 mM KCl, 5 mM EDTA, 10% Glycerol, and 1 mM PMSF), followed by incubation with 15 μl of protein G-conjugated Sepharose beads prebound with 1 μl anti-FLAG antibody. Samples were slowly rotated at 4°C overnight, followed by washing (three times) with 1 ml of buffer (0.05% digitonin, 25 mM Tris-HCl, pH 7.5, 50 mM KCl, 5 mM EDTA, 10% Glycerol and 1 mM PMSF). The immunoprecipitated proteins were heated at 55°C with 1X SDS buffer, eluted by running the samples in SDS-PAGE, and analysed by western blotting using the respective antibodies.

### Mass spectrometry analysis for protein identification

The supernatant fraction obtained from co-immunoprecipitation was mixed with 0.1 ml urea buffer (8 M), which was further mixed with 5 µl of 200 mM DTT in 50 mM ammonium bicarbonate and incubated at room temperature for 1h. Next, 20 µl of 200 mM iodoacetamide dissolved in 50 mM ammonium bicarbonate was added to the samples. The mixture was then incubated at room temperature for 1h in the dark, followed by adding 50 µl of 1 mM CaCl_2_ in 50 mM ammonium bicarbonate to the reaction solution and incubated for 10 min at room temperature. For protease digestion, trypsin was added at a 1:30 (trypsin:protein) ratio, and the samples were desalted using a C18 column with 50 µl of 50 mM ammonium bicarbonate and 0.1% trifluoroacetic acid (pH 3-4). The C18 column was activated using 200 µl of 50% acetonitrile (activation solution), centrifuged at 1500 g for 1 min, and then treated with 200 µl of 0.5% trifluoroacetic acid in 20% acetonitrile. The mixed samples were loaded into the column multiple times, and short spins discarded the flow-through. 200 µl 0.5% trifluoroacetic acid in 5% acetonitrile was also used to wash the column by passing multiple times. The fragmented peptides (bound to the column) were eluted using 50 µl of 0.1% trifluoroacetic acid in 70% acetonitrile. The eluted fraction was dried and resuspended in 0.1% trifluoroacetic acid containing 2% acetonitrile, and the sample was measured. LC-MS/MS on an UltiMate™ 3000 Peptide analysis system was performed on a Fusion mass spectrometer (Thermo Fisher Scientific) coupled with an RSLCnano system. A 110 min gradient was employed for peptide separation. Electrospray and negative spray voltages of 2200 V and 600 V, respectively, were used. Subsequent tandem mass spectrometry (MS/MS) analysis was carried out to identify peptides. Intact peptides were detected in the Orbitrap mass analyser at a resolution of 60,000. Peptide ions were selected for fragmentation using a normalised collision energy (NCE) setting of 30. Fragment ions were detected in the ion trap mass analyser using centroid mode. The resulting peptide sequence data were searched against the UniProt database using Proteome Discoverer 2.5.

### Analysis of mitochondrial morphology

The deletion and mutant strains were transformed with pRS415_TEF_ MTS-mCherry and pRS416_TEF_ MTS-mCherry vectors to visualise mitochondrial morphology, respectively. MTS-mCherry specifically decorates mitochondria, enabling visualisation of mitochondrial morphology(Bankapalli *et al*, 2015). The cells were harvested at mid-log phase (OD_600_ ∼0.5) and subjected to acetic acid stress. The harvested cells were then washed once with 1X PBS, spread onto agarose pads, and visualised under a DeltaVision Elite Fluorescence Microscope (GE Healthcare) using a 100X objective lens. The λ_EX_/λ_EM_ for mCherry used was 587/610 nm.

### Flow cytometry experiments

The DNA content profile in cells was determined using propidium iodide (PI) staining, as described previously with slight modifications(Riccardi & Nicoletti, 2006). The deletion and mutant strains cultured in SC Dextrose media were pelleted and fixed in 70% ethanol overnight at −20°C. The fixed cells were washed with 1X PBS and centrifuged at 2000 rpm for 5 min. Cells were then resuspended in 0.2 ml of 1X PBS and 0.2 ml of DNA extraction buffer (1.92 ml of 0.2 M Na_2_HPO_4_ with 0.8 ml of 0.1% Triton X-100 (v/v), pH 7.8) and incubated for 5 min at room temperature. Cells were centrifuged and dissolved in 0.2 ml of DNA staining solution (1 mg/ml of PI in 1 ml of PBS with 2 mg of DNase-free RNase). The resuspended cells were incubated for 30 min at room temperature in the dark, then subjected to flow cytometry. BD FACS Verse Flow Cytometer (BD Biosciences) with a 488 nm excitation laser was used to analyse 30,000 cells per experiment at a low flow rate. FlowJo 10.0 software was used for data analysis.

To measure total and functional mitochondrial mass, WT, deletion, and mutant strains were grown to early log phase (OD_600_ ∼0.2-0.3) and subjected to acetic acid stress. The cells were then washed once with 1X PBS, followed by staining with 10 µM NAO (Nonyl Acridine Orange) for 20 min and 8.75 µM TMRE (Tetramethyl rhodamine ethyl ester) for 15 min in the dark at 30°C to detect total and functional mitochondrial mass, respectively. The cells were then pelleted, dissolved in 1X PBS and subjected to flow cytometry analysis on a BD FACSVerse™ flow cytometer. The excitation/emission wavelengths used for NAO and TMRE were 488/520 nm and 549/575 nm, respectively. 10,000 events were recorded per sample, and the data were analysed using FlowJo 10.0. For both total and functional mitochondrial mass, relative median fluorescence values were represented as bar graphs by quantifying the fold change in median fluorescence intensity of the deletion and mutant strains with respect to the wild-type control.

### Measurement of ROS levels and mtDNA assessment

The WT, deletion, and mutant strains were grown to the early log phase (OD_600_ ∼0.5), subjected to 70 mM acetic acid stress, and then incubated with 5 μM MitoSox dye for 30 min in the dark at 30°C. The harvested cells were divided and viewed under a Floview 3000 Olympus confocal microscope with a 100X objective lens and analysed using CellSens software after deconvolution. The remaining cells were used for FACS analysis on a BD FACSVerse™ flow cytometer. The excitation/emission wavelengths used for MitoSox are 510/580 nm.

Similarly, the mtDNA analysis method was followed with modifications(Hull *et al*, 2008; Susarla *et al*., 2023). The above-mentioned yeast cells were treated with acetic acid, harvested, and stained with 10 μM SYTO18 for 15 min at 30°C. The harvested cells were washed once with 1X PBS, then spread onto agarose pads and visualised under a Floview 3000 Olympus confocal microscope using a 100X objective lens. The λ_EX_/λ_EM_ for mtDNA was 488/533 nm.

### Statistical analysis

Protein band intensities were quantified using MultiGauge software. Subsequent data analysis and statistical tests were performed in Excel (Microsoft) and GraphPad Prism 8.2.4, respectively. Statistical analyses were performed using unpaired one-way and two-way analysis of variance (ANOVA) (with multiple comparisons). All quantitative data were obtained from at least 2 independent experiments, and error bars represent the standard error of the mean (SEM). Asterisks denote the following levels of significance: *, *p* ≤ 0.05; **, *p* ≤ 0.01; ***, *p* ≤ 0.001; ****, *p* ≤ 0.0001.

## Author Contributions

A.C., R.D., S.S., and P.D.S designed and conceptualised the project. A.C. designed and carried out the genetics, biochemical, and proteomics studies. R.D. performed and analysed the microscopic and FACS data. A.C., R.D., S.S., and P.D.S interpreted data, and A.C. and R.D. wrote the manuscript. S.S. and P.D.S. edited and reviewed the manuscript.

## Acknowledgements

This work was supported by the Anusandhan National Research Foundation (ANRF) (ANRF/ARG/2025/001223/LS) and the Department of Science and Technology (DST-FIST Program-Phase III, no. SR/FST/LSII-045/2016-G). Aishita Chakraborty and Dr. Rachayeeta Deb acknowledge the fellowship from the Department of Biotechnology as a Ph.D. scholar (DBT-JRF/SRF) and research associate (DBT-RA), India, respectively. We want to extend our gratitude to Prof. Dr. Johannes Herrmann (University of Kaiserslautern, Germany) for gifting the anti-Nde1 antibody. We sincerely acknowledge Dr. SreeDivya Saladi for her critical discussion and valuable feedback for this study. We gratefully thank Dr. Abhishek Kumar for providing the Tim22 clone constructs required for these studies. We acknowledge the Department of Biochemistry for the Confocal Imaging and BD Cytoflex FACS facility and the Division of Biological Sciences, Indian Institute of Science, for the Mass Spectrometry and Flow Cytometry facility.

## References

Baker MJ, Frazier AE, Gulbis JM, Ryan MT (2007) Mitochondrial protein-import machinery: correlating structure with function. Trends in Cell Biology 17: 456–464

Bankapalli K, Saladi S, Awadia SS, Goswami AV, Samaddar M, D’Silva P (2015) Robust Glyoxalase activity of Hsp31, a ThiJ/DJ-1/PfpI Family Member Protein, Is Critical for Oxidative Stress Resistance in Saccharomyces cerevisiae. Journal of Biological Chemistry 290: 26491–26507

Bankapalli K, Vishwanathan V, Susarla G, Sunayana N, Saladi S, Peethambaram D, D’Silva P (2020) Redox-dependent regulation of mitochondrial dynamics by DJ-1 paralogs in Saccharomyces cerevisiae. Redox Biology 32: 101451

Büttner S, Eisenberg T, Herker E, Carmona-Gutierrez D, Kroemer G, Madeo F (2006) Why yeast cells can undergo apoptosis: death in times of peace, love, and war. The Journal of Cell Biology 175: 521–525

Callegari S, Richter F, Chojnacka K, Jans DC, Lorenzi I, Pacheu-Grau D, Jakobs S, Lenz C, Urlaub H, Dudek J et al (2016) TIM29 is a subunit of the human carrier translocase required for protein transport. FEBS Letters 590: 4147–4158

Chacinska A, Koehler CM, Milenkovic D, Lithgow T, Pfanner N (2009) Importing Mitochondrial Proteins: Machineries and Mechanisms. Cell 138: 628–644

Chacinska A, Lind M, Frazier AE, Dudek J, Meisinger C, Geissler A, Sickmann A, Meyer HE, Truscott KN, Guiard B et al (2005) Mitochondrial Presequence Translocase: Switching between TOM Tethering and Motor Recruitment Involves Tim21 and Tim17. Cell 120: 817–829

Chaves SR, Rego A, Martins VM, Santos-Pereira C, Sousa MJ, Côrte-Real M (2021) Regulation of Cell Death Induced by Acetic Acid in Yeasts. Frontiers in Cell and Developmental Biology 9

Chiusolo V, Jacquemin G, Yonca Bassoy E, Vinet L, Liguori L, Walch M, Kozjak-Pavlovic V, Martinvalet D (2017) Granzyme B enters the mitochondria in a Sam50-, Tim22- and mtHsp70-dependent manner to induce apoptosis. Cell Death & Differentiation 24: 747–758

Ciesielski GL, Plotka M, Manicki M, Schilke BA, Dutkiewicz R, Sahi C, Marszalek J, Craig EA (2013) Nucleoid localization of Hsp40 Mdj1 is important for its function in maintenance of mitochondrial DNA. Biochim Biophys Acta 1833: 2233–2243

Clifford J, Chiba H, Sobieszczuk D, Metzger D, Chambon P (1996) RXRalpha-null F9 embryonal carcinoma cells are resistant to the differentiation, anti-proliferative and apoptotic effects of retinoids. The EMBO Journal 15: 4142–4155

D’Silva PR, Schilke B, Hayashi M, Craig EA, Brodsky J (2008) Interaction of the J-Protein Heterodimer Pam18/Pam16 of the Mitochondrial Import Motor with the Translocon of the Inner Membrane. Molecular Biology of the Cell 19: 424–432

Dadsena S, Jenner A, García-Sáez AJ (2021) Mitochondrial outer membrane permeabilization at the single molecule level. Cellular and Molecular Life Sciences 78: 3777–3790

Daniel Gietz R, Woods RA (2002) Transformation of yeast by lithium acetate/single-stranded carrier DNA/polyethylene glycol method. In: *Guide to Yeast Genetics and Molecular and Cell Biology - Part B*, pp. 87–96.

Del Carratore R, Della Croce C, Simili M, Taccini E, Scavuzzo M, Sbrana S (2002) Cell cycle and morphological alterations as indicative of apoptosis promoted by UV irradiation in S. cerevisiae. Mutation Research/Genetic Toxicology and Environmental Mutagenesis 513: 183–191

Du L, Yu Y, Li Z, Chen J, Liu Y, Xia Y, Liu X (2007) Tim18, a component of the mitochondrial translocator, mediates yeast cell death induced by arsenic. Biochemistry (Moscow) 72: 843–847

Dudek J, Rehling P, van der Laan M (2013) Mitochondrial protein import: Common principles and physiological networks. Biochimica et Biophysica Acta (BBA) - Molecular Cell Research 1833: 274–285

Falcone C, Mazzoni C (2016) External and internal triggers of cell death in yeast. Cellular and Molecular Life Sciences 73: 2237–2250

Gebert N, Chacinska A, Wagner K, Guiard B, Koehler CM, Rehling P, Pfanner N, Wiedemann N (2008) Assembly of the three small Tim proteins precedes docking to the mitochondrial carrier translocase. EMBO reports 9: 548–554

Gebert N, Gebert M, Oeljeklaus S, von der Malsburg K, Stroud David A, Kulawiak B, Wirth C, Zahedi René P, Dolezal P, Wiese S et al (2011) Dual Function of Sdh3 in the Respiratory Chain and TIM22 Protein Translocase of the Mitochondrial Inner Membrane. Molecular Cell 44: 811–818

Gentle IE, Perry AJ, Alcock FH, Likic VA, Dolezal P, Ng ET, Purcell AW, McConnville M, Naderer T, Chanez AL et al (2007) Conserved Motifs Reveal Details of Ancestry and Structure in the Small TIM Chaperones of the Mitochondrial Intermembrane Space. Molecular Biology and Evolution 24: 1149–1160

Giannattasio S, Atlante A, Antonacci L, Guaragnella N, Lattanzio P, Passarella S, Marra E (2008) Cytochrome c is released from coupled mitochondria of yeast en route to acetic acid-induced programmed cell death and can work as an electron donor and a ROS scavenger. FEBS Letters 582: 1519–1525

Giannattasio S, Guaragnella N, Ždralević M, Marra E (2013) Molecular mechanisms of Saccharomyces cerevisiae stress adaptation and programmed cell death in response to acetic acid. Frontiers in Microbiology 4

Grandier-Vazeille X, Bathany K, Chaignepain S, Camougrand N, Manon S, Schmitter J-M (2001) Yeast Mitochondrial Dehydrogenases Are Associated in a Supramolecular Complex. Biochemistry 40: 9758–9769

Green DR, Narendra DP, Jin SM, Tanaka A, Suen D-F, Gautier CA, Shen J, Cookson MR, Youle RJ (2010) PINK1 Is Selectively Stabilized on Impaired Mitochondria to Activate Parkin. PLoS Biology 8

Guaragnella N, Bettiga M (2021) Acetic acid stress in budding yeast: From molecular mechanisms to applications. Yeast 38: 391–400

Guo Y, Cheong N, Zhang Z, De Rose R, Deng Y, Farber SA, Fernandes-Alnemri T, Alnemri ES (2004) Tim50, a Component of the Mitochondrial Translocator, Regulates Mitochondrial Integrity and Cell Death. Journal of Biological Chemistry 279: 24813–24825

Harbauer Angelika B, Zahedi René P, Sickmann A, Pfanner N, Meisinger C (2014) The Protein Import Machinery of Mitochondria—A Regulatory Hub in Metabolism, Stress, and Disease. Cell Metabolism 19: 357–372

Herrmann JM, Riemer J (2021) Apoptosis inducing factor and mitochondrial NADH dehydrogenases: redox-controlled gear boxes to switch between mitochondrial biogenesis and cell death. Biological Chemistry 402: 289–297

Hull CM, Ingavale SS, Chang YC, Lee H, McClelland CM, Leong ML, Kwon-Chung KJ (2008) Importance of Mitochondria in Survival of Cryptococcus neoformans Under Low Oxygen Conditions and Tolerance to Cobalt Chloride. PLoS Pathogens 4

Jackson TD, Hock DH, Fujihara KM, Palmer CS, Frazier AE, Low YC, Kang Y, Ang C-S, Clemons NJ, Thorburn DR et al (2021) The TIM22 complex mediates the import of sideroflexins and is required for efficient mitochondrial one-carbon metabolism. Molecular Biology of the Cell 32: 475–491

Ježek J, Cooper K, Strich R (2018) Reactive Oxygen Species and Mitochondrial Dynamics: The Yin and Yang of Mitochondrial Dysfunction and Cancer Progression. Antioxidants 7

Kalpage HA, Wan J, Morse PT, Zurek MP, Turner AA, Khobeir A, Yazdi N, Hakim L, Liu J, Vaishnav A et al (2020) Cytochrome c phosphorylation: Control of mitochondrial electron transport chain flux and apoptosis. The International Journal of Biochemistry & Cell Biology 121

Kang Y, Anderson AJ, Jackson TD, Palmer CS, De Souza DP, Fujihara KM, Stait T, Frazier AE, Clemons NJ, Tull D et al (2019) Function of hTim8a in complex IV assembly in neuronal cells provides insight into pathomechanism underlying Mohr-Tranebjærg syndrome. eLife 8

Kang Y, Baker MJ, Liem M, Louber J, McKenzie M, Atukorala I, Ang C-S, Keerthikumar S, Mathivanan S, Stojanovski D (2016) Tim29 is a novel subunit of the human TIM22 translocase and is involved in complex assembly and stability. eLife 5

Kang Y, Fielden LF, Stojanovski D (2018) Mitochondrial protein transport in health and disease. Seminars in Cell & Developmental Biology 76: 142–153

Kang Y, Stroud DA, Baker MJ, De Souza DP, Frazier AE, Liem M, Tull D, Mathivanan S, McConville MJ, Thorburn DR et al (2017) Sengers Syndrome-Associated Mitochondrial Acylglycerol Kinase Is a Subunit of the Human TIM22 Protein Import Complex. Molecular Cell 67: 457–470.e455

Kasahara A, Scorrano L (2014) Mitochondria: from cell death executioners to regulators of cell differentiation. Trends in Cell Biology 24: 761–770

Kerscher O, Sepuri NB, Jensen RE, Fox TD (2000) Tim18p Is a New Component of the Tim54p-Tim22p Translocon in the Mitochondrial Inner Membrane. Molecular Biology of the Cell 11: 103–116

Kizmaz B, Nutz A, Egeler A, Herrmann JM (2024) Protein insertion into the inner membrane of mitochondria: routes and mechanisms. FEBS Open Bio 14: 1627–1639

Koehler CM (2004) New Developments in Mitochondrial Assembly. Annual Review of Cell and Developmental Biology 20: 309–335

Koehler CM, Murphy MP, Bally NA, Leuenberger D, Oppliger W, Dolfini L, Junne T, Schatz G, Or E (2000) Tim18p, a New Subunit of the TIM22 Complex That Mediates Insertion of Imported Proteins into the Yeast Mitochondrial Inner Membrane. Molecular and Cellular Biology 20: 1187–1193

Kovermann P, Truscott KN, Guiard B, Rehling P, Sepuri NB, Müller H, Jensen RE, Wagner R, Pfanner N (2002) Tim22, the Essential Core of the Mitochondrial Protein Insertion Complex, Forms a Voltage-Activated and Signal-Gated Channel. Molecular Cell 9: 363–373

Kumar A, Matta SK, D’Silva P (2020) Conserved regions of budding yeast Tim22 have a role in structural organization of the carrier translocase. Journal of Cell Science 133: jcs244632

Kumar A, Waingankar TP, D’Silva P (2023) Functional crosstalk between the TIM22 complex and YME1 machinery maintains mitochondrial proteostasis and integrity. Journal of Cell Science 136

Lalier L, Mignard V, Joalland M-P, Lanoé D, Cartron P-F, Manon S, Vallette FM (2021) TOM20-mediated transfer of Bcl2 from ER to MAM and mitochondria upon induction of apoptosis. Cell Death & Disease 12

Li W, Sun L, Liang Q, Wang J, Mo W, Zhou B (2006) Yeast AMID Homologue Ndi1p Displays Respiration-restricted Apoptotic Activity and Is Involved in Chronological Aging. Molecular Biology of the Cell 17: 1802–1811

Lionaki E, Gkikas I, Tavernarakis N (2023) Mitochondrial protein import machinery conveys stress signals to the cytosol and beyond. BioEssays 45

Liu Q, D’Silva P, Walter W, Marszalek J, Craig EA (2003) Regulated Cycling of Mitochondrial Hsp70 at the Protein Import Channel. Science 300: 139–141

Ludovico P, Sousa MJ, Silva MT, Leão Cl, Côrte-Real M (2001) Saccharomyces cerevisiae commits to a programmed cell death process in response to acetic acid. Microbiology 147: 2409–2415

Luttik MAH, Overkamp KM, Kötter P, de Vries S, van Dijken JP, Pronk JT (1998) The Saccharomyces cerevisiae NDE1 and NDE2 Genes Encode Separate Mitochondrial NADH Dehydrogenases Catalyzing the Oxidation of Cytosolic NADH. Journal of Biological Chemistry 273: 24529–24534

Madeo F, Carmona-Gutierrez D, Ring J, Büttner S, Eisenberg T, Kroemer G (2009) Caspase-dependent and caspase-independent cell death pathways in yeast. Biochemical and Biophysical Research Communications 382: 227–231

Matta SK, Pareek G, Bankapalli K, Oblesha A, D’Silva P (2017) Role of Tim17 Transmembrane Regions in Regulating the Architecture of Presequence Translocase and Mitochondrial DNA Stability. Molecular and Cellular Biology 37

Meisinger C, Pfanner N, Truscott KN (2006) Isolation of yeast mitochondria. Methods Mol Biol 313: 33–39

Mokranjac D, Popov-Čeleketić D, Hell K, Neupert W (2005) Role of Tim21 in Mitochondrial Translocation Contact Sites *. Journal of Biological Chemistry 280: 23437–23440

Mussulini BHM, Maruszczak KK, Draczkowski P, Borrero-Landazabal MA, Ayyamperumal S, Wnorowski A, Wasilewski M, Chacinska A (2025) MIA40 suppresses cell death induced by apoptosis-inducing factor 1. *EMBO Reports*

Mustafa M, Ahmad R, Tantry IQ, Ahmad W, Siddiqui S, Alam M, Abbas K, Moinuddin, Hassan MI, Habib S et al (2024) Apoptosis: A Comprehensive Overview of Signaling Pathways, Morphological Changes, and Physiological Significance and Therapeutic Implications. Cells 13

Neupert W, Herrmann JM (2007) Translocation of Proteins into Mitochondria. Annual Review of Biochemistry 76: 723–749

Newton K, Strasser A, Kayagaki N, Dixit VM (2024) Cell death. Cell 187: 235–256

Nunnari J, Suomalainen A (2012) Mitochondria: In Sickness and in Health. Cell 148: 1145–1159

O’Malley J, Kumar R, Inigo J, Yadava N, Chandra D (2020) Mitochondrial Stress Response and Cancer. Trends in Cancer 6: 688–701

Okamoto H, Miyagawa A, Shiota T, Tamura Y, Endo T (2014) Intramolecular Disulfide Bond of Tim22 Protein Maintains Integrity of the TIM22 Complex in the Mitochondrial Inner Membrane. Journal of Biological Chemistry 289: 4827–4838

Oliveira CSF, Pereira H, Alves S, Castro L, Baltazar F, Chaves SR, Preto A, Côrte-Real M (2015) Cathepsin D protects colorectal cancer cells from acetate-induced apoptosis through autophagy-independent degradation of damaged mitochondria. Cell Death & Disease 6: e1788–e1788

Palmer CS, Anderson AJ, Stojanovski D (2021) Mitochondrial protein import dysfunction: mitochondrial disease, neurodegenerative disease and cancer. FEBS Letters 595: 1107–1131

Pareek G, Krishnamoorthy V, D’Silva P (2013) Molecular Insights Revealing Interaction of Tim23 and Channel Subunits of Presequence Translocase. Molecular and Cellular Biology 33: 4641–4659

Pfanner N, Warscheid B, Wiedemann N (2019) Mitochondrial proteins: from biogenesis to functional networks. Nature Reviews Molecular Cell Biology 20: 267–284

Qi L, Wang Q, Guan Z, Wu Y, Shen C, Hong S, Cao J, Zhang X, Yan C, Yin P (2020) Cryo-EM structure of the human mitochondrial translocase TIM22 complex. Cell Research 31: 369–372

Rehling P, Brandner K, Pfanner N (2004) Mitochondrial import and the twin-pore translocase. Nature Reviews Molecular Cell Biology 5: 519–530

Rehling P, Model K, Brandner K, Kovermann P, Sickmann A, Meyer HE, Kühlbrandt W, Wagner R, Truscott KN, Pfanner N (2003) Protein Insertion into the Mitochondrial Inner Membrane by a Twin-Pore Translocase. Science 299: 1747–1751

Riccardi C, Nicoletti I (2006) Analysis of apoptosis by propidium iodide staining and flow cytometry. Nature Protocols 1: 1458–1461

Roesch K (2002) Human deafness dystonia syndrome is caused by a defect in assembly of the DDP1/TIMM8a-TIMM13 complex. Human Molecular Genetics 11: 477–486

Saladi S, Boos F, Poglitsch M, Meyer H, Sommer F, Mühlhaus T, Schroda M, Schuldiner M, Madeo F, Herrmann JM (2020) The NADH Dehydrogenase Nde1 Executes Cell Death after Integrating Signals from Metabolism and Proteostasis on the Mitochondrial Surface. Molecular Cell 77: 189–202.e186

San-Millán I (2023) The Key Role of Mitochondrial Function in Health and Disease. Antioxidants 12

Sirrenberg C, Bauer MF, Guiard B, Neupert W, Brunner M (1996) Import of carrier proteins into the mitochondrial inner membrane mediated by Tim22. Nature 384: 582–585

Small WC, McAlister-Henn L (1998) Identification of a Cytosolically Directed NADH Dehydrogenase in Mitochondria of Saccharomyces cerevisiae. Journal of Bacteriology 180: 4051–4055

Susarla G, Kataria P, Kundu A, D’Silva P (2023) Saccharomyces cerevisiae DJ-1 paralogs maintain genome integrity through glycation repair of nucleic acids and proteins. eLife 12

van der Laan M, Hutu DP, Rehling P (2010) On the mechanism of preprotein import by the mitochondrial presequence translocase. Biochimica et Biophysica Acta (BBA) - Molecular Cell Research 1803: 732–739

Vukotic M, Nolte H, König T, Saita S, Ananjew M, Krüger M, Tatsuta T, Langer T (2017) Acylglycerol Kinase Mutated in Sengers Syndrome Is a Subunit of the TIM22 Protein Translocase in Mitochondria. Molecular Cell 67: 471–483.e477

Wang X (2001) The expanding role of mitochondria in apoptosis. Genes Dev 15: 2922–2933

Webb CT, Gorman MA, Lazarou M, Ryan MT, Gulbis JM (2006) Crystal Structure of the Mitochondrial Chaperone TIM9•10 Reveals a Six-Bladed α-Propeller. Molecular Cell 21: 123–133

Weiss F, Lauffenburger D, Friedl P (2022) Towards targeting of shared mechanisms of cancer metastasis and therapy resistance. Nature Reviews Cancer 22: 157–173

Wiedemann N, Pfanner N (2017) Mitochondrial Machineries for Protein Import and Assembly. Annu Rev Biochem 86: 685–714

Wittig I, Braun H-P, Schägger H (2006) Blue native PAGE. Nature Protocols 1: 418–428

Wrobel L, Sokol AM, Chojnacka M, Chacinska A (2016) The presence of disulfide bonds reveals an evolutionarily conserved mechanism involved in mitochondrial protein translocase assembly. Scientific Reports 6

Wrobel L, Topf U, Bragoszewski P, Wiese S, Sztolsztener ME, Oeljeklaus S, Varabyova A, Lirski M, Chroscicki P, Mroczek S et al (2015) Mistargeted mitochondrial proteins activate a proteostatic response in the cytosol. Nature 524: 485–488

Zorov DB, Juhaszova M, Sollott SJ (2014) Mitochondrial Reactive Oxygen Species (ROS) and ROS-Induced ROS Release. Physiological Reviews 94: 909–950

